# Identification of genomic features that uniquely impact estrogen receptor alpha binding and its effects on gene expression in endometrial cancer

**DOI:** 10.64898/2025.12.11.693723

**Authors:** Alexandra Richey, Noel Kitchen, Craig M. Rush, Jeffery M. Vahrenkamp, Hosiana Abewe, Kyle S. Osbourne, Alex J. Bott, Brian K. Dalley, Jared Rutter, Jason Gertz

## Abstract

Estrogen receptor alpha (ER) is an established oncogenic transcription factor in breast and endometrial cancer; however, more is known about the mechanisms controlling ER behavior in breast cancer, and therapies targeting ER have been much more successful in breast cancer. To address this disparity, we characterize the genomic features that control ER in endometrial cancer and determine to what extent these factors differ from those in breast cancer. We focus on the locations of estrogen response elements (EREs), ER’s preferred DNA binding motif, throughout the human genome. To identify factors that predict ER genomic binding and effects on target gene expression, we apply machine learning to genomic data for each ERE in Ishikawa cells (ER-positive endometrial cancer) and T-47D cells (ER-positive breast cancer). Many of these factors, such as chromatin accessibility and histone modifications, are predictive of ER activity in both cell lines. However, the transcription factors that predict ER activity are cell type-specific, including FOXA1 and GATA3 in T-47D cells, and ETV4 and SOX17 in Ishikawa cells. In addition, the features that predict ER binding and effects on gene expression differ, with transcription at EREs in the absence of estrogen being predictive of ER regulatory activity. A CRISPR knockout screen in Ishikawa cells, and follow-up experiments, confirm the discovery that SOX17 controls ER activity in endometrial cancer cells. These results identify important genomic features of ER binding and regulatory activity and how these features differ between endometrial cancer and breast cancer cells.

## Introduction

Estrogen receptor alpha (ER) is an established oncogene in the hormone-dependent forms of breast and endometrial cancers (Jordan and Brodie 2007; Paterni et al. 2014; Rodriguez et al. 2019). While ER signaling has been extensively studied in breast cancer, leading to the development of targeted therapies such as selective estrogen receptor modulators (SERMs) and degraders (SERDs) (Jordan 2006; Lee et al. 2017), similar therapeutic advancements have not been achieved in endometrial cancer. For instance, tamoxifen—a SERM that functions as an ER antagonist in breast tissue—exerts agonistic effects in the endometrium, preventing its use as an endometrial cancer treatment (Hu et al. 2015). Despite its established clinical significance, ER and the related molecular mechanisms contributing to endometrial cancer progression remain poorly understood. With the incidence and mortality of endometrial cancer rising year after year (Siegel et al. 2025), there is an urgent need to understand the molecular mechanisms of endometrial cancer in order to identify therapeutic targets and improve patient outcomes.

One likely contributor to the limited translatability of the ER-targeting therapies of breast cancer to endometrial cancer is the tissue type-specific behavior of ER. Less than 10% of ER genomic binding sites are shared between the Ishikawa (ER+ endometrial cancer) and the T-47D (ER+ breast cancer) cell lines (Gertz et al. 2013), and ER binding site patterns in patient samples segregate distinctly by tumor type (Droog et al. 2016; Rodriguez et al. 2020). Cofactors of ER also differ between breast and endometrial cancer. FOXA1 and GATA3 are essential for regulating ER binding in the breast cancer context (Hurtado et al. 2011; Kong et al. 2011; Theodorou et al. 2013), and their corresponding motifs are uniquely enriched at ER binding sites in breast cancer patient samples compared to endometrial cancer patient samples (Ross-Innes et al. 2012; Rodriguez et al. 2020). However, GATA3 is rarely expressed in endometrial cancer, and FOXA1 has minimal overlap with ER genomic binding in endometrial cancer (Droog et al. 2016; Rodriguez et al. 2020). ETV4 has been shown to regulate ER binding in endometrial cancer cells, but not in breast cancer cells, and the ETS and SOX motifs are uniquely enriched at ER binding sites in endometrial cancer patient samples compared to breast cancer patient samples (Rodriguez et al. 2020).

The preferred DNA binding motif of ER is the estrogen response element (ERE) (Klein-Hitpass et al. 1988), which is important for recruiting ER to gene expression enhancers (Carleton et al. 2020). Like other transcription factors, ER does not bind to all EREs within the genome (Gertz et al. 2013) and disruption of ER-bound EREs does not always alter gene expression (Carleton et al. 2017; Ginley-Hidinger et al. 2019; Abewe et al. 2025). This suggests that the presence of an ERE alone is insufficient to predict ER occupancy or transcriptional regulatory activity. The locations of EREs across the genome are consistent across breast and endometrial cancer cells, providing the opportunity to make direct comparisons of their regulatory potential between tissue types.

Machine learning approaches provide a powerful tool for characterizing the ER-mediated molecular mechanisms in endometrial cancer and uncovering how these mechanisms differ from those of breast cancer. Many large-scale datasets have been generated for breast and endometrial cancer (Ginley-Hidinger et al. 2024; Abewe et al. 2025), and machine learning allows us to uncover important features of potential regulatory regions. Gregoricchio et al. successfully used machine learning to identify an enhancer mutation that alleviates transcriptional repression of ER during endometrial tumorigenesis (Gregoricchio et al. 2025). In our previous work, we successfully used Boruta (Kursa and Rudnicki 2010), a feature selection algorithm, to identify predictors of timing, noise, and expression levels of the transcriptional response to estrogen (Ginley-Hidinger et al. 2024) and to identify predictors of changes in 3D genomic interactions in response to estrogen (Abewe et al. 2025). Other researchers have also used Boruta to identify gene variants that may lead to disease states (Ragunthar and Selvakumar 2019). Here, we apply Boruta to the study of EREs that bind ER and impact gene expression. We also performed a CRISPR knockout (KO) screen of epigenetic and transcription factors to identify important contributors to ER activity in endometrial cancer cells.

## Results

### Very few estrogen response elements in the genome are bound by ER and affect gene expression

To evaluate the usage of EREs by ER, we first identified 82,534 significant full ERE matches in the human female genome (see Methods). We considered a bound ERE to be one that overlapped with an ER ChIP-seq peak from the corresponding cell line (Fig. 1A). We considered an ERE that affects gene expression, termed “regulatory,” to be one that is located near the transcription start site (TSS) of a gene that changes expression in response to estrogen, using a 100kb distance cutoff based on our previous analysis (Fig. 1A) (Carleton et al. 2017). In Ishikawa cells, 2154 (2.6%) EREs are bound by ER, and 935 (43.4% of bound EREs) EREs affect expression (Fig. 1B). In T-47D cells, 2329 (2.8%) EREs are bound, and 1005 (43.2% of bound EREs) EREs affect expression (Fig. 1B). In Ishikawa cells, approximately 20% of regulatory EREs are associated with both up- and downregulated genes, 62% are only associated with upregulated genes, and 18% are only associated with downregulated genes. In T-47D cells, approximately 12% of regulatory EREs are associated with both up- and downregulated genes, 67% are only associated with upregulated genes, and 21% are only associated with downregulated genes. Ishikawa and T-47D cells have been shown to have roughly equal levels of ER expression, which matches their similar levels of bound and regulatory EREs (Gertz et al. 2013). Between the cell lines, 895 of the bound EREs (24.9% of all bound EREs) and 335 of the regulatory EREs (20.09% of all regulatory EREs) are shared in both cell lines. With the vast majority of EREs not bound by ER in a given cell line, we evaluated whether the number of EREs in the human genome was mostly due to random chance or if there were signs of evolutionary pressure to maintain a high number of EREs in the genome. We counted the EREs on Chromosome 1 and then shuffled the sequence of Chromosome 1 ten times and recounted the EREs. Chromosome 1 contains 6922 EREs compared to an average number of 5005 EREs in the shuffled chromosomes, with a standard deviation of 87. Therefore, the number of EREs on Chromosome 1 is 22 standard deviations higher than the mean from the shuffled chromosomes (Fig. 1C), suggesting that there is some evolutionary pressure to preserve the high number of EREs in the human genome, even if most remain unbound by ER within a cell type.

**Figure 1:**
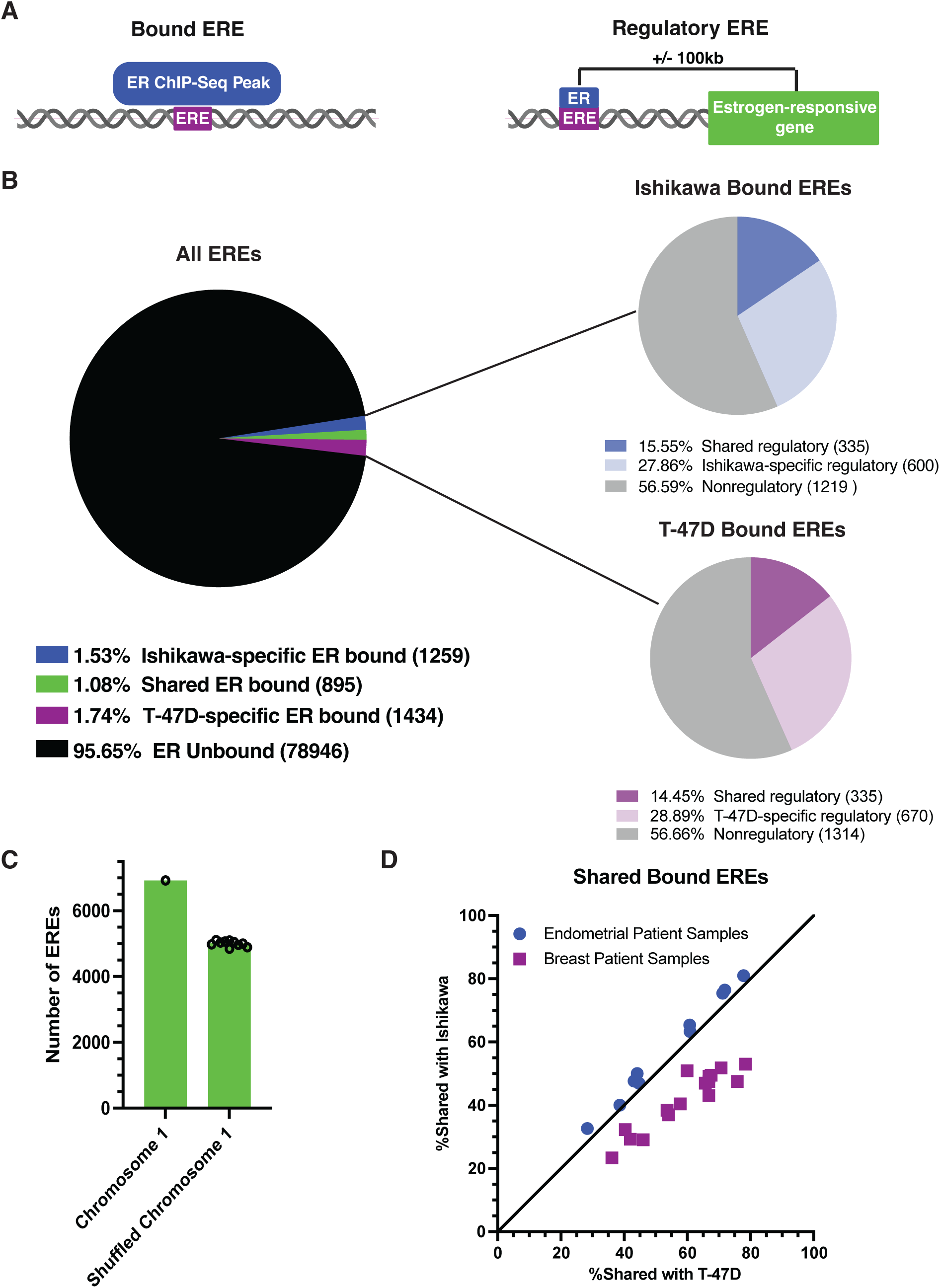
Most EREs are unbound by ER and do not affect gene expression. A) Illustration shows the definition of bound EREs (left) and regulatory EREs (right). B) Pie chart on the left shows the number of EREs that are bound in both cell lines, bound only in Ishikawa, bound only in T-47D, and unbound. Pie charts on the right show, out of the EREs that are bound in Ishikawa (top) and T-47D (bottom), the number of regulatory EREs unique to that cell line, regulatory EREs shared between cell lines, and nonregulatory EREs. C) Bar chart shows the number of EREs found in Chromosome 1, compared to the number of EREs in ten different random shufflings of Chromosome 1. D) Scatter plot displays the percentage of bound EREs shared with the Ishikawa cell line (y-axis) and with the T-47D cell line (x-axis) from endometrial cancer patient samples (blue circles) and breast cancer patient samples (purple squares). All patient samples share more bound EREs with their corresponding cell line than with their non-corresponding cell line.

Analysis of ER ChIP-seq data from breast and endometrial cancer patient samples (Rodriguez et al. 2020) was used to determine how many EREs were bound in the patient samples, and how many bound EREs they shared with the Ishikawa and T-47D cell lines (Fig. 1D). The average percentage of bound EREs in endometrial cancer patient samples was 1.63% with a standard deviation of 1.22%, and the average percentage of bound EREs in breast cancer patient samples was 1.59% with a standard deviation of 1.36%. The bound EREs in endometrial cancer patient samples matched more closely with the bound EREs in the Ishikawa cell line (Wilcoxon signed-rank test; *P*-value = 9.766×10^-4^), and the bound EREs in breast cancer patient samples matched more closely with the bound EREs in the T-47D cell line (Wilcoxon signed-rank test, *P*-value = 1.526×10^-5^). For our further analyses, we focus on Ishikawa and T-47D cell lines, since they represent the broad differences observed in ER binding between endometrial and breast cancer specimens.

### Several factors predict ERE binding in a cell type-specific manner

To identify factors that can predict whether an ERE will be bound by ER we compiled the following datasets: the relative affinity of the ERE for ER (determined by the sequence of the ERE, see Methods), ChIP-seq for multiple transcription factors and histone modifications, PRO-seq, DNase-seq, ATAC-seq, and HiChIP to determine the number of 3D connections associated with each ERE (Table S1). These datasets were previously published, with the exception of the SOX17 ChIP-seq data in Ishikawa cells. We previously found that the SOX motif is enriched at ER binding sites in endometrial cancer specifically (Rodriguez et al. 2020), and *SOX17* is the most highly expressed SOX factor in Ishikawa cells. Our datasets differed somewhat between Ishikawa cells and T-47D cells due to differences in data availability. We focused only on datasets taken from cells grown in the absence of estrogen, to ensure that the data represented a state preceding ER binding, rather than a state caused by ER binding.

In order to identify which factors are predictive of an ERE’s ability to bind ER, we used the Boruta feature selection algorithm (Kursa and Rudnicki 2010). In the Ishikawa cells, 22 factors were in the initial dataset tested in Boruta. Of those, 19 were confirmed important in predicting ER binding, 1 was tentative, and 2 were rejected (Fig. 2A, Table S2). In the T-47D cells, 21 factors were in the initial dataset tested in Boruta. Of those, 18 were confirmed important in predicting ER binding, and 3 were rejected (Fig. 2B, Table S2). For all factors in both cell lines that were confirmed as important, their average value was higher for bound EREs than for unbound EREs, suggesting that all predictors are positively associated with ER binding (Fig. 2A, B), which is consistent with a focus on factors related to active gene regulation. Because there is a large imbalance between the number of bound and unbound EREs, we also performed 100 Boruta iterations on subsamples of our dataset with equal numbers of bound and unbound EREs. These results largely aligned with the results from running Boruta on the full dataset, with a few factors, such as CTCF, changing their designation (Supp. Fig. 1, Table S2).

**Figure 2:**
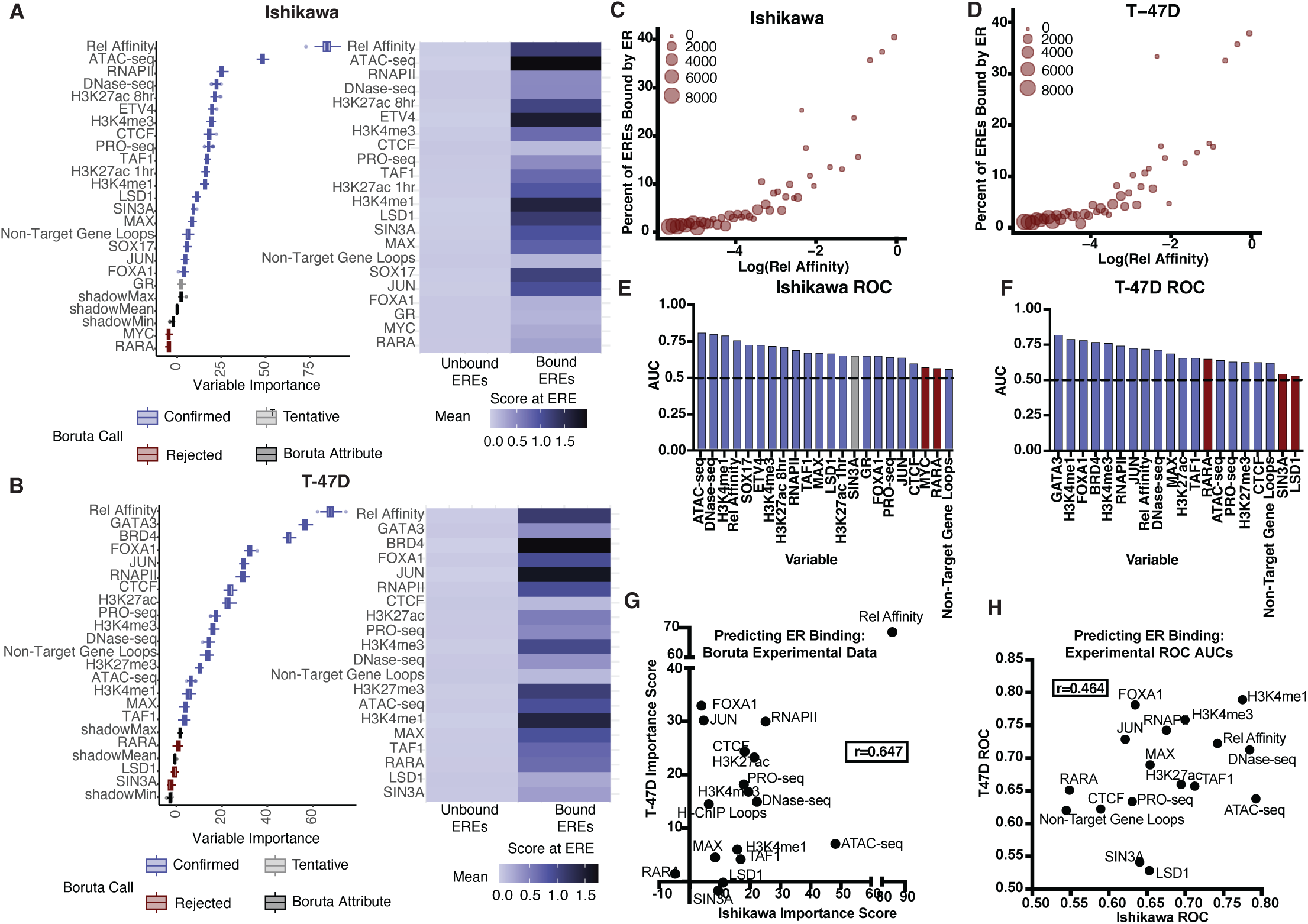
Bound EREs are predicted by sequence, chromatin accessibility, histone modifications, and cell line-specific transcription factors. Boxplots show the ranked factor importance for predicting bound EREs by Boruta for A) (left) Ishikawa and B) (left) T-47D. Colors indicate Boruta determination of importance; blue is confirmed, light gray is tentative, red is rejected, and dark gray is a Boruta attribute. Heatmaps show the normalized average value of each factor at bound and unbound EREs for A) (right) Ishikawa and B) (right) T-47D. Scatter plots show the relationship between the log of Relative Affinity for ER of an ERE and the percentage of EREs bound by ER for C) Ishikawa and D) T-47D. Bar charts show the ROC AUC of each factor individually in predicting bound EREs in E) Ishikawa and F) T-47D. Bar colors correspond with Boruta determination of importance. The dotted line shows the value 0.5, indicating an ROC AUC from random prediction. G) Scatter plot shows the comparison of Boruta importance scores in predicting bound EREs between cell lines from the experimental data, with a Pearson correlation coefficient of 0.647. H) Scatter plots show the comparison of ROC AUCs in predicting bound EREs between cell lines from the experimental dataset, with a Pearson correlation coefficient of 0.464.

For both cell lines, the top scoring factor in predicting ER binding was relative affinity for ER (Fig. 2A, B). Further analysis showed that relative affinity is positively correlated with the percentage of EREs bound by ER (Fig. 2C, D), indicating that the closer an ERE sequence is to the optimal ERE, the more likely it is to be bound. Relative affinity is higher on average in bound EREs shared between the cell lines than in EREs that are uniquely bound in one cell line or the other (Wilcoxon rank-sum test, *P*-value =1.2116×10^-42^). Despite the high importance of relative affinity, optimal EREs, EREs with the highest possible relative affinity, were only bound by ER 40% of the time, indicating that ER binding affinity is mostly determined by other factors. For optimal EREs, 46% are in open chromatin in Ishikawa cells, and 11% are in T-47D cells. Of these optimal EREs in open chromatin, 60% are bound by ER in Ishikawa cells, and 79% in T-47D cells. The percentage of EREs in open chromatin bound by ER drops as relative affinity drops. For EREs scoring in the lowest 10% of relative affinity, 21% are in open chromatin in Ishikawa cells, and 6% in T-47D cells. Of these low-scoring EREs in open chromatin, 3% are bound in Ishikawa cells, and 8% in T-47D cells.

Additional factors found to be predictive in both cell lines included chromatin accessibility as measured by DNase-seq and ATAC-seq, and activating histone modifications, including H3K4me1. For T-47D cells, FOXA1 and GATA3 binding were found to be predictive, which supports the validity of Boruta, as FOXA1 and GATA3 have been established to control ER binding in the context of breast cancer (Hurtado et al. 2011; Kong et al. 2011; Theodorou et al. 2013). ETV4, which we previously identified as a contributor to ER binding in endometrial cancer (Rodriguez et al. 2020), was identified as predictive of ER binding in the Ishikawa cells (Fig. 2A). The Boruta results are supported by the receiver operator characteristic curve (ROC) analysis, where factors mentioned above exhibited areas under the curve (AUC) substantially greater than 0.5 (Fig. 2E, F).

### Motif analysis reveals more cell line-specific predictors of ER binding

In order to expand our analysis beyond experimentally assayed transcription factors, we used the HOMER motif database (Heinz et al. 2010), which provides the genomic locations of the cognate DNA binding motifs for 450 DNA-binding proteins. We first performed clustering of the 450 motifs to remove redundancy, which created 270 motif clusters (Table S3), and then performed Boruta on these 270 motif clusters. For this dataset, each factor was binary: either an ERE had one of the motifs in the cluster within 100bp of the ERE, or it did not.

In Ishikawa cells, 103 clusters were confirmed as important in predicting ER binding, with 32 clusters being designated as tentative, and 135 clusters being rejected as unimportant in predicting ER binding (Table S3). In T-47D cells, 98 clusters were confirmed as important in predicting ER binding, with 26 clusters being designated as tentative, and 146 clusters being rejected in predicting ER binding (Table S3). However, many of these confirmed motif clusters contained motifs that are very similar to the ERE motif. Therefore, we removed any motifs containing at least an ERE half site from further investigation. To narrow down our focus, we further analyzed two groups of clusters: 1) clusters that had an importance score higher than 10, and 2) clusters that were confirmed in only one cell line.

All the motif clusters that had an importance score above 10 in either cell line were confirmed as important for predicting binding in both cell lines. Eight clusters in Ishikawa and 12 clusters in T-47D had an importance score over 10 (Supp. Fig. 2A). We observed some notable motifs in the clusters that were confirmed in only one cell line (Supp. Fig. 2B, C). The compound motif of NF1:FOXA1 was confirmed only in T-47D, along with a cluster containing the GATA3 motif. The cluster containing the SOX17 motif was only confirmed in Ishikawa. These results support our findings from experimental datasets, while identifying some additional factors that may contribute to ER binding.

### Factors that contribute to ER binding are mostly cell type-specific

For both cell lines, relative affinity for ER had the highest Boruta importance score (Fig. 2G). It also had high predictive ability in ROC AUC analysis in both cell lines. The relative affinity of ERE is the exact same in both cell lines; therefore, ERE strength doesn’t explain why different sets of EREs are bound in the two cell lines. Of the genomic features assayed in both cell lines, we found several factors that are better at predicting ER binding in one cell line over the other. In both Boruta and ROC AUC analyses, FOXA1 and JUN were better predictors in T-47D cells, while ATAC-seq, LSD1, and SIN3A had higher predictive ability in Ishikawa cells (Fig. 2G, H). Most motif clusters had similarly poor predictive ability in both cell lines, but there were a handful of outliers for one cell line or the other (Supp. Fig. 2D). In addition to the features assessed in both cell lines, transcription factors that were predicted to be important for ER binding, GATA3, ETV4, and SOX17, were only examined in one cell line because of cell type-specific expression (Supp. Fig. 3A) (Gertz et al. 2013). The cell type-specific expression of these factors, as well as FOXA1, is also observed in patient tumor data from The Cancer Genome Atlas (TCGA) (Supp Fig. 3B-E) (The ICGC/TCGA Pan-Cancer Analysis of Whole Genomes Consortium 2020). Overall, these results are consistent with previous findings that ER binding is controlled by different factors in breast and endometrial cancer cells.

**Figure 3:**
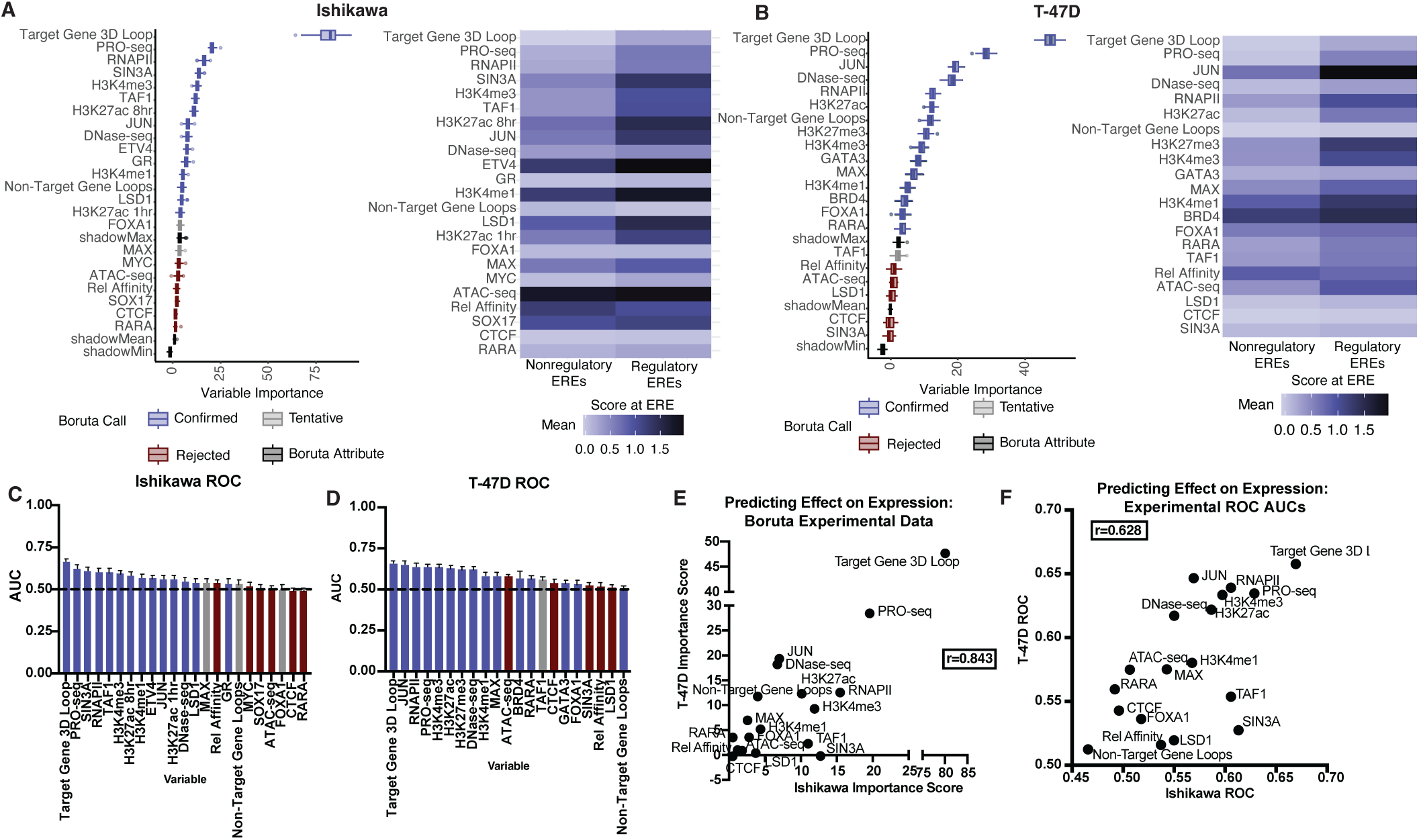
Regulatory EREs are enriched for 3D genomic interactions and active transcription. Boxplots show the ranked factor importance for predicting regulatory EREs by Boruta for A) (left) Ishikawa and B) (left) T-47D. Colors indicate Boruta determination of importance; blue is confirmed, light gray is tentative, red is rejected, and dark gray is a Boruta attribute. Heatmaps show the normalized average value of each factor at regulatory and nonregulatory EREs for A) (right) Ishikawa and B) (right) T-47D. Bar charts show the ROC AUC of each factor individually in predicting regulatory EREs in C) Ishikawa and D) T-47D. Bar colors correspond to Boruta determination of importance. The dotted line shows the value 0.5, indicating an ROC AUC from random prediction. E) Scatter plot shows the comparison of Boruta importance scores in predicting regulatory EREs between cell lines from the experimental data, with a Pearson correlation coefficient of 0.843. F) Scatter plot show the comparison of ROC AUCs in predicting regulatory EREs between cell lines from the experimental dataset, with a Pearson correlation coefficient of 0.628.

### Predictive multi-variable model of ERE binding

With the results from Boruta and the ROC AUC analyses, we sought to create multivariable logistic regression models that could predict, with high sensitivity and specificity, whether any given ERE would be bound by ER. For both cell lines, we ran two-variable logistic regression for all possible combinations using our non-motif dataset, using down-sampling while creating the training groups to account for the unbalanced dataset. We then ranked the variables based on the average ROC AUC across their two-variable logistic regression models (Supp. Fig. 4A, B). These ranks correlated well with the Boruta importance score ranks in both cell lines, with Pearson correlation coefficients of 0.671 and 0.592 for Ishikawa and T-47D, respectively. As a baseline for more complex models, we started with the single variable relative affinity because of its high Boruta importance score and because it performed well in all two-variable analyses. For Ishikawa EREs, this model can predict which EREs will be bound with a sensitivity of 63.5% and a specificity of 75.9% (ROC AUC = 0.741, Supp. Fig. 4C). For T-47D EREs, the model has a sensitivity of 60.4% and a specificity of 74.9% (ROC AUC = 0.720, Supp. Fig. 4D). These models can be improved upon, as relative affinity is the same between the cell lines, and a different set of EREs is bound in each cell line. To do this, we added ATAC-seq, a measure of chromatin accessibility, and ChIP-seq of H3K4me1, a histone modification associated with active enhancers. Even though these datasets are predictive of ER binding in both cell lines, they can explain the cell line differences, as the genomic locations of accessible chromatin and H3K4me1 differ between T-47D and Ishikawa cells (Gertz et al. 2013). For Ishikawa cells, adding ATAC-seq and H3K4me1 ChIP-seq improved the sensitivity to 78.5%, the specificity to 87.4%, and the ROC AUC to 0.907 (Supp. Fig. 4C). For T-47D cells, it improved the sensitivity to 73.0%, the specificity to 84.9%, and the ROC AUC to 0.892 (Supp. Fig. 4D). Another way to improve the model is adding cell type-specific transcription factors. For Ishikawa cells, having the variables of relative affinity, ETV4 ChIP-seq, and SOX17 ChIP-seq creates a model with 75.1% sensitivity, 83.5% specificity, and an ROC AUC of 0.869 (Supp. Fig. 4C). For T-47D cells, having the variables of relative affinity, FOXA1 ChIP-seq and GATA3 ChIP-seq creates a model with 75.6% sensitivity, 87.1% specificity, and an ROC AUC of 0.893 (Supp. Fig. 4D). These results suggest that the sequence of the ERE, accessible chromatin, and histone modifications associated with active enhancer activity, such as H3K4me1, are important determinants of ER binding potential, regardless of cell line. However, sequence-specific transcription factors that determine ER binding potential are cell line-specific, with FOXA1 and GATA3 determining ER binding potential in T-47D cells and ETV4 and SOX17 determining ER binding potential in Ishikawa cells.

**Figure 4:**
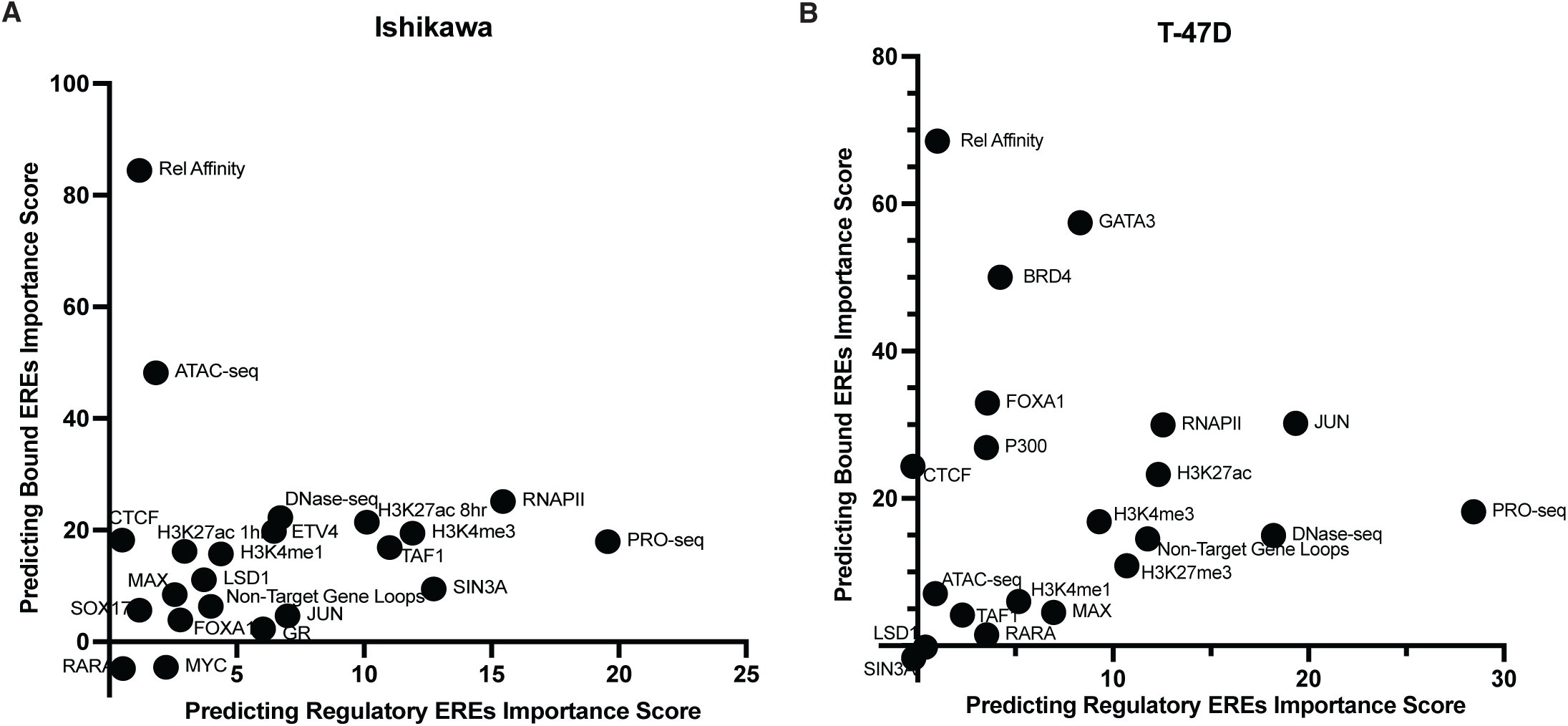
Different factors are predictive of bound EREs and regulatory EREs. Scatter plots show Boruta importance scores of each factor in experimental data for predicting bound EREs vs. predicting regulatory EREs in A) Ishikawa and B) T-47D.

To validate our findings in another endometrial cancer model, we performed ATAC-seq, H3K4me1 ChIP-seq, and SOX17 ChIP-seq in HCI-EC-23 (Rush et al. 2022) to recreate our logistic regression models for predicting ER binding. We were unable to perform ETV4 ChIP-seq for technical reasons. HCI-EC-23 has lower ER activity than Ishikawa cells, as reflected by it having only 386 bound EREs. Relative affinity alone has a sensitivity of 63.4% and a specificity of 75.9% (ROC AUC=0.742, Supp. Fig. 4E). Adding SOX17 increased the sensitivity to 70.7% and the specificity to 80.3% (ROC AUC=0.833, Supp. Fig. 4E). When we added ATAC-seq and H3K4me1 ChIP-seq to relative affinity, the model improved to a sensitivity of 78.0%, a specificity of 87.5%, and an ROC AUC of 0.905. A model including all four variables has a sensitivity of 78.8%, a specificity of 87.7%, and an ROC AUC of 0.908 (Supp. Fig. 4E). The performance of these models is very similar to Ishikawa cells, suggesting that the same features are important for ERE binding across endometrial cancer models.

The models are only slightly improved by combining all five variables. For Ishikawa cells, a model with all five variables has a 79.5% sensitivity, 87.9% specificity, and an ROC AUC of 0.913 (Supp. Fig. 4C). For T-47D, a model with all five variables has a 79.4% sensitivity, 88.4% specificity, and an ROC AUC of 0.913 (Supp. Fig. 4D). Similarly, when all possible variables are used, the Ishikawa model has an 80.4% sensitivity, 88.0% specificity, and an ROC AUC of 0.917, and the T-47D model has an 80.5% sensitivity, 90.0% specificity, and an ROC AUC of 0.924.

### Transcription and chromatin looping predict regulatory EREs

To identify factors that are predictive of a bound ERE’s ability to affect gene expression, we analyzed the same datasets as were used when analyzing binding. We also added the factor “Target Gene 3D Loop,” which is a binary classifier that indicates whether an ERE has a 3D genomic interaction with the transcription start site of an estrogen-regulated gene, based on HiChIP data. We first compared all regulatory EREs against all nonregulatory bound EREs. For the Ishikawa cells, 15 factors were confirmed important in predicting whether an ERE would affect gene expression, 2 were tentative, and 6 were rejected (Fig. 3A, Table S2). For T-47D cells, 16 factors were confirmed, 1 was tentative, and 5 were rejected (Fig. 3B, Table S2). For both cell lines, the top two scoring features were Target Gene 3D Loop and PRO-seq. This suggests that 3D genomic interactions and active transcription at promoter distal EREs in the absence of estrogen, as measured by PRO-seq, are both important in distinguishing regulatory EREs from nonregulatory EREs. Once again, the average value of each factor confirmed important by Boruta was higher for regulatory EREs than for nonregulatory EREs, indicating that our dataset is biased toward factors that promote ER activity (Fig. 3A, B). We also performed Boruta analysis comparing only promoter-distal regulatory EREs to nonregulatory EREs, only regulatory EREs of upregulated genes to nonregulatory EREs, and only regulatory EREs of downregulated genes to nonregulatory EREs. The results from these analyses were very similar to the analyses including all regulatory EREs (Table S4 and Supp. Fig. 5).

**Figure 5:**
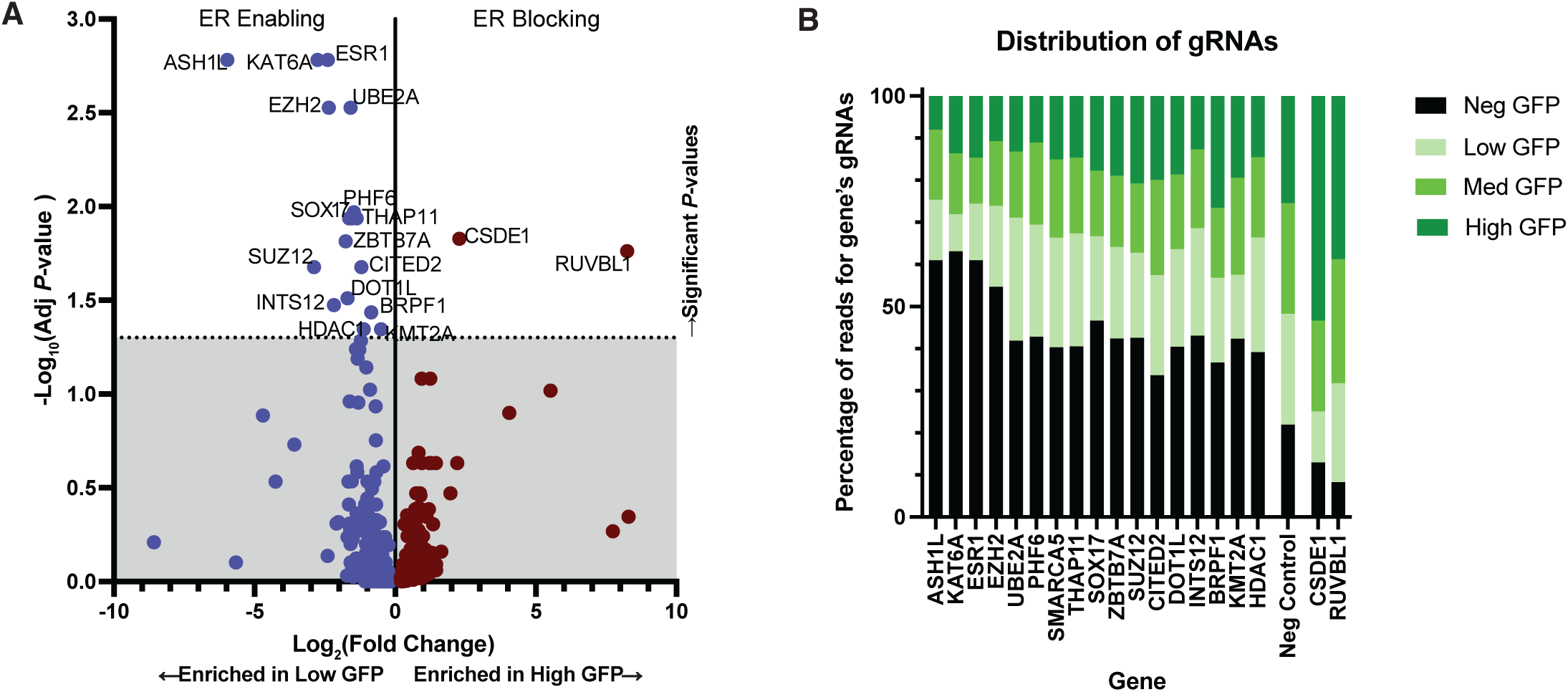
CRISPR KO screen identifies important genes for ER activity in Ishikawa cells. A) Volcano plot shows the ER enabling and ER blocking genes with their log_2_ fold changes and adjusted *P*-values. B) Stacked bar plot shows the gRNA counts for each significant gene in the four GFP bins. The gRNAs for the ER enabling genes are enriched in the negative GFP bin (black), and the gRNAs for the ER blocking genes are enriched in the high GFP bin (darkest green).

We also calculated ROC AUCs for this dataset, and Target Gene 3D Loop and PRO-seq were both highly ranked in Ishikawa and T-47D. However, overall, the ROC AUCs for predicting regulatory EREs were lower than those for predicting bound EREs (Fig. 3C, D). In Ishikawa cells, the average ROC AUC for predicting bound EREs was 0.663, and the average ROC AUC for predicting regulatory EREs was 0.557 (Wilcoxon rank-sum test, *P*-value = 4.62×10^-7^). In T-47D cells, the average ROC AUC for predicting bound EREs was 0.689, and the average ROC AUC for predicting regulatory EREs was 0.579 (Wilcoxon rank-sum test, *P*-value = 4.31×10^-6^).

We next determined which motifs are predictive of an ERE’s ability to affect gene expression. Very few of the 270 motif clusters were found to be important; 9 were confirmed important in the Ishikawa dataset, and 11 in the T-47D dataset, while 7 were tentative for Ishikawa, and 3 for T-47D (Supp. Fig. 6A, B, Table S3). Only one cluster was confirmed in both cell lines: the Maz motif, which is very GC rich. It should also be noted that relative affinity for ER scored very poorly for predicting regulatory EREs (Fig. 3A, B). In general, sequence-based features are less predictive of regulatory EREs than experimental datasets such as ChIP-seq, PRO-seq, and HiChIP.

**Figure 6:**
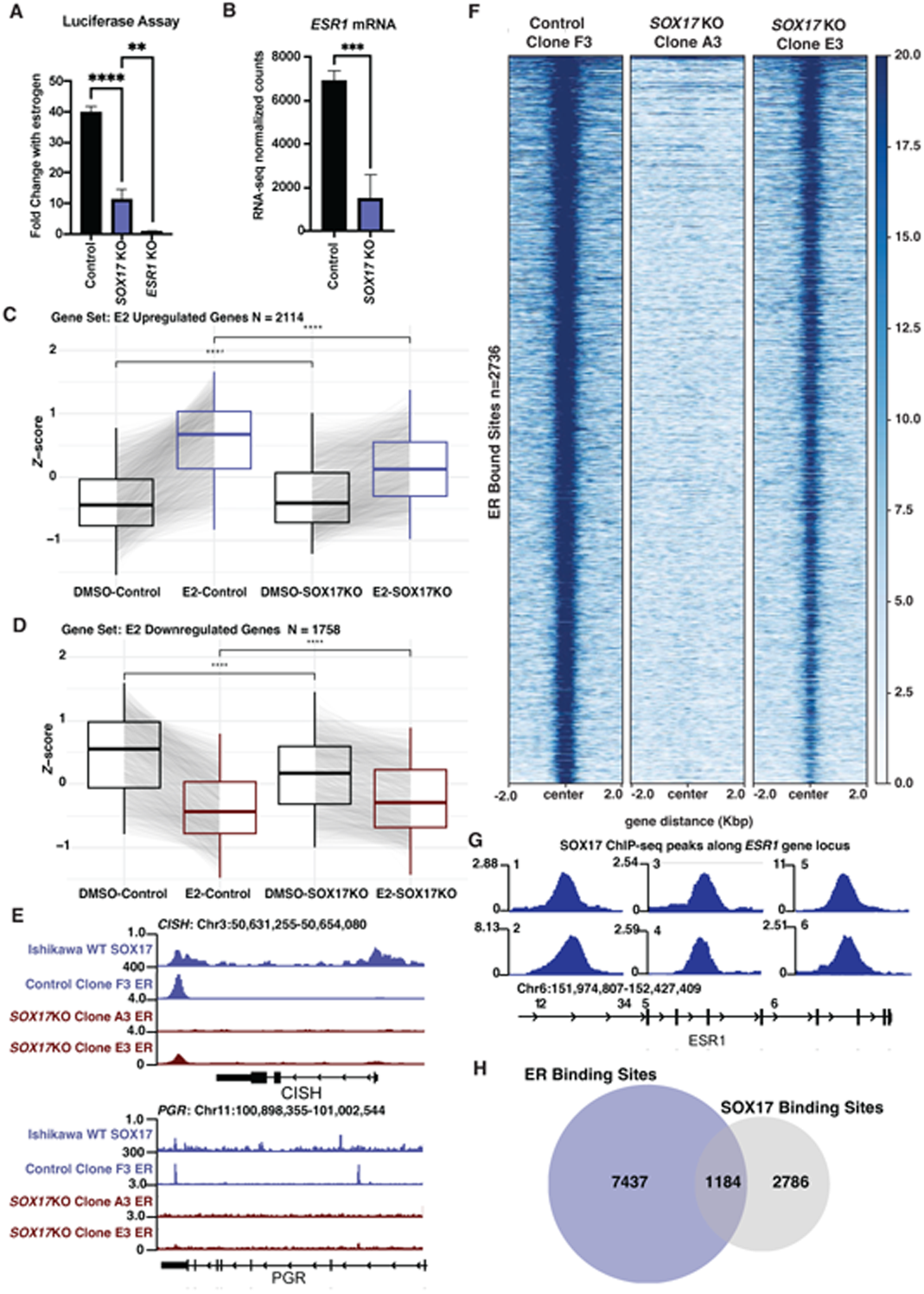
SOX17 KO decreases ER activity. A) Bar chart shows ERE luciferase reporter activity relative to no estrogen treatment in SOX17 KO lines compared to a control and an ER KO line. B) Bar graph shows decreased *ESR1* mRNA normalized counts in SOX17 KO lines compared to control. Aggregate plots show *Z*-scores comparing DMSO treated and estrogen (E2) treated samples from the RNA-seq data for SOX17 KO lines and control lines for C) E2 upregulated genes and D) E2 downregulated genes. E) Browser track shows decreased ER binding at estrogen responsive genes *CISH* and *PGR.* F) Heatmap shows decreased ER genomic binding signal in SOX17 KO lines compared to control lines across all ER bound sites. G) Browser track shows six SOX17 binding sites near *ESR1* gene locus. H) Venn diagram shows overlap of binding sites between ER and SOX17 in parental Ishikawa cells.

### Similar features predict regulatory EREs across cell lines

When we directly compare Ishikawa and T-47D regulatory ERE associated factors, we observed several similarities. For both cell lines, interacting with an estrogen responsive gene via looping and transcription prior to estrogen treatment at the location of the ERE are the top predictors of a bound ERE’s ability to affect gene expression. This is shown both with Boruta importance scores and ROC AUCs (Fig. 3E, F). We observed high correlation coefficients in both Boruta importance scores (r=0.843) and ROC AUCs (r=0.628). JUN ChIP-seq is a better predictor in T-47D cells, and SIN3A ChIP-seq is a better predictor in Ishikawa cells, just as they were for predicting binding. Supplemental Figure 6C shows the comparisons of importance scores from the Boruta analysis using motif data. This data shows very little concordance between the cell lines; however, since motif data overall was not found to be very effective in predicting regulatory EREs, the lack of correlation is expected.

### Predictive multi-variable model of regulatory EREs

As we did with ERE binding, we sought to create a multivariable logistic regression model that could identify regulatory EREs with high sensitivity and specificity. We performed logistic regression using all possible two-variable combinations from the non-motif datasets and ranked variables by their mean ROC AUC (Supp. Fig. 7A, B). As with ER binding, these ranks correlated well with the Boruta importance score ranks in both cell lines, with Pearson correlation coefficients of 0.872 and 0.671 for Ishikawa and T-47D, respectively. As a baseline, we used the single variable model for Target Gene 3D Loop, since it had the highest importance score in both cell lines in the Boruta analysis and performed well in all two-variable models. For Ishikawa cells, that model has a sensitivity of 34.8%, a specificity of 99.1%, and an ROC AUC of 0.670 (Supp. Fig. 7C). For T-47D cells, that model has a sensitivity of 42.0%, a specificity of 89.9%, and an ROC AUC of 0.659 (Supp. Fig. 7D). To expand the model for Ishikawa cells, we used Target Gene 3D Loop, PRO-seq, and SIN3A ChIP-seq, which returned a sensitivity of 40.1%, a specificity of 94.9%, and an ROC AUC of 0.743 (Supp. Fig. 7C). For the T-47D multi-variable model, we used Target Gene 3D Loop, PRO-seq, and JUN ChIP-seq, which returned a sensitivity of 44.4%, a specificity of 88.7%, and an ROC AUC of 0.721 (Supp. Fig. 7D). From these models and the results of the Boruta analyses, we can conclude that 3D genomic interactions and active transcription at promoter distal enhancers are important determinants of a bound ERE’s regulatory potential, along with the presence of SIN3A in Ishikawa cells and the presence of JUN in T-47D cells. However, our models achieved high specificity only and had low sensitivity, indicating that these factors can accurately identify a regulatory ERE, but most regulatory EREs are missed.

### Different factors associate with bound EREs and regulatory EREs

The factors that predict whether an ERE will bind ER and the factors that predict whether a bound ERE will affect gene expression are largely different. In both Ishikawa and T-47D cells, relative affinity is only predictive of whether an ERE will bind ER, and PRO-seq is more predictive of whether a bound ERE will affect gene expression than it is of whether an ERE will bind ER (Fig. 4A, B). Differences between the cell lines are more prevalent in predicting binding than in predicting effects on expression. There is higher correlation between the Boruta importance scores shown in the scatter plot for predicting regulatory EREs (r=0.843, Fig. 3E) than there is shown in the scatter plot for predicting bound EREs (r=0.647, Fig. 2G). There are strong differences in estrogen-mediated gene regulation between breast cancer cells and endometrial cancer cells, and these results suggest that these differences arise more prominently from where ER binds within the genome than from how ER regulates gene expression once bound to an ERE.

### CRISPR KO screen reveals factors important for ER activity in Ishikawa cells

In order to functionally evaluate the importance of different factors in mediating ER activity, we performed a CRISPR KO screen targeting 2500 genes encoding epigenetic and transcription factors that could influence ER activity. To measure ER activity, we created an Ishikawa line with an integrated GFP reporter driven by three consecutive EREs. The reporter line was transduced with the Human Epigenetic Knockout lentiviral library (Williams et al. 2020). After selection, growth, and 72 hours of estrogen treatment, cells were sorted into four bins based on their GFP levels (negative, low, medium, and high). Genes whose guide RNAs were enriched in the negative GFP bin compared to the other three bins were determined to be ER enabling genes, and genes whose guide RNAs were enriched in the high GFP bin compared to the other three bins were determined to be ER blocking genes (Table S5). We identified 18 significant genes (Fig. 5), including *ESR1*, the gene encoding ER. In general, the ER enabling genes supported conclusions made from the Boruta and ROC AUC analyses. As expected, there were several significant genes encoding histone modifying enzymes, including *ASH1l, KAT6A, KMT2A,* and *BRPF1*, all of which are associated with transcriptional activation. Two ER blocking genes were identified: *CSDE1* and *RUVBL1*. CSDE1 has been shown to both enable and block gene expression in the cancer context as an RNA-binding protein (Ciocia et al. 2024). *RUVBL1* encodes an H4 and H2A acetyltransferase, which is typically associated with transcriptional activation (Doyon et al. 2004); therefore, it may indirectly block ER activity.

### SOX17 is important for ER activity in Ishikawa cells

*SOX17* was identified as an ER enabling gene in the CRISPR KO screen, which agrees with our computational analyses, suggesting that SOX17 regulates ER activity in endometrial cancer. In support of this role, *SOX17* is more highly expressed in endometrial tumor tissue than in normal tissue or breast cancer (Supp. Fig. 3E). *SOX17* expression is also positively correlated with *ESR1* expression in endometrial cancer, with a Spearman correlation of 0.556 (Supp. Fig. 8A). Based on DepMap data (Arafeh et al. 2025), endometrial cancer models are more dependent on *SOX17* compared to breast cancer models (Wilcoxon signed-rank test, *P*-value = 6.5×10^-4^) and models from other cancer types (Wilcoxon signed-rank test, *P*-value = 0.0017), with the exception of ovarian cancer models (Supp. Fig. 8B). Together, our results suggest that SOX17 could be an important factor in promoting ER activity in endometrial cancer.

To further investigate the role of SOX17 in regulating ER activity in endometrial cancer cells, we created *SOX17* KO lines derived from Ishikawa cells using CRISPR and individually cloned guide RNAs. We also created *ESR1* KO Ishikawa cells as a positive control. We verified the absence of SOX17 and ER protein, in the respective KOs (Supp. Fig. 9A-E). To assess effects on ER activity, we performed an ERE reporter assay, similar to the one used in the CRISPR screen, and found that ER activity was decreased from 40-fold, over no estrogen controls, to 10-fold in *SOX17* KO lines*; ESR1* KO drove ERE levels back to no estrogen control levels as expected (Fig. 6A). To investigate the effects on estrogen-mediated gene expression, we performed RNA-seq with and without estrogen treatment in *SOX17* KO lines and control lines. *ESR1* expression itself was affected, decreasing from an average of 6899 normalized RNA-seq counts in control lines to an average of 1472 normalized counts in *SOX17* KO lines (Fig. 6B). In control lines, the expression of 3835 genes was significantly impacted by estrogen treatment, whereas only 317 genes changed expression in response to estrogen treatment in the *SOX17* KO lines. The magnitude of expression responses, for both upregulated and downregulated genes, was significantly reduced in *SOX17* KO cells (Fig. 6C, D, Supp. Fig. 9F).

ER genomic binding was also reduced in *SOX17* KO lines at estrogen responsive genes such as *CISH* and *PGR* (Fig. 6E), and across the genome (Fig. 6F). ER ChIP-seq in *SOX17* KO lines produced an average of 429 peaks compared to 2550 peaks in wild-type cells (Fig. 6F). The reduction in ER activity may be due to two separate causes: 1) SOX17 regulates ER transcription by binding to six sites near the *ESR1* gene locus (Fig. 6G), consistent with the 5-fold decrease in *ESR1* expression levels in the *SOX17* KO cells (Fig. 6B); and 2) SOX17 binding overlaps with ER at many genomic sites, potentially aiding in ER-driven transcription regulation. In parental Ishikawa cells, ER has 8621 binding sites in the presence of estrogen, SOX17 has 3970 in the absence of estrogen, and they overlap at 1184 sites (30% of SOX17 bound sites; Fig. 6H), suggesting that SOX17 binding at these sites may help facilitate ER binding. Overall, we found that the presence of SOX17 significantly affects ER protein levels, activity, and genomic binding in endometrial cancer cells.

## Discussion

Estrogen signaling, working through ER, plays an oncogenic role in breast and endometrial cancer. However, the genomic binding sites and target genes of ER differ significantly between the cancer types. To explore the causes of cell type specificity, we decided to use EREs as a consistent anchor between breast and endometrial cancer cells. In cell lines and tumor samples from both cancer types, very few EREs (less than 3%) contribute to genomic binding or gene regulation. Of the bound EREs or regulatory EREs, about a quarter are shared between breast cancer and endometrial cancer cells. Despite the low level of ERE utilization, there are significantly more EREs in the human genome than can be expected by random chance, suggesting that EREs are under evolutionary pressure to be conserved. It is possible that these EREs are involved in gene regulation in other tissues or in different phases of development. The highly selective usage of EREs provided an opportunity to evaluate the factors that contribute to ER genomic binding and gene regulation and how they differ between cell types.

To identify important features of EREs that bind ER in endometrial cancer and breast cancer cells, we compiled several datasets in Ishikawa and T-47D cells. Boruta, which assigns importance scores to features based on a random forest algorithm, revealed several factors that associate with ER binding. The highest scoring factor for both Ishikawa and T-47D cells was relative affinity, meaning the closer an ERE’s sequence is to the optimal ER binding sequence, the more likely it is to be bound by ER. However, only 40% of EREs with the highest possible relative affinity are bound, suggesting other features are key in controlling ER binding. Other features found to be enriched at bound EREs in both the Ishikawa and T-47D lines were features correlating with accessible chromatin and histone modifications associated with active enhancer regions, both of which are found at active transcription factor binding sites (Field and Adelman 2020; Panigrahi and O’Malley 2021). Patterns of open chromatin are cell type-specific (Gertz et al. 2013), supporting the cell type specificity of ER binding, but these chromatin patterns are controlled by sequence specific transcription factors.

Our analysis uncovered several transcription factors associated with bound EREs, many of which were expected based on previous studies of ER control in breast and endometrial cancer. FOXA1 and GATA3 were both found to be highly predictive of bound EREs in the T-47D breast cancer cell line. FOXA1 and GATA3 act as pioneer factors and facilitate ER binding in the breast cancer context by associating with chromatin ahead of ER recruitment (Hurtado et al. 2011; Theodorou et al. 2013; Jiang et al. 2019). ETV4 was found to be predictive of ER binding in the Ishikawa endometrial cancer cell line, which is supported by our previous work showing that ETV4 is essential for ER activity in endometrial cancer cells by binding with ER at many of its genomic sites and helping to localize ER to the nucleus (Rodriguez et al. 2020). With these results, we built both a transcription factor focused model using cell type-specific transcription factors and a chromatin focused model using ATAC-seq and H3K4me1; each could predict binding with reasonable sensitivity and specificity. For endometrial cancer, we were able to verify the predictive power of our models in a second cell line. Even though HCI-EC-23 has less ER activity than Ishikawa, and less bound EREs, we were still able to construct a model that predicted bound EREs with similar sensitivity and specificity to the Ishikawa models for predicting ERE binding.

To expand the search beyond previous experimental data, we analyzed motif instances, and the results generally supported our findings from the experimental data. The FOXA1-containing motif cluster was more predictive of ER binding in T-47D than Ishikawa cells, the GATA3 motif cluster was predictive of ER binding in T-47D cells only, and the SOX17 motif cluster was predictive of ER binding in Ishikawa cells only. Other findings from the motif analyses warrant further investigation. For example, the ZIC2, GLIS3, and various KLF motifs scored highly in both cell lines. ZIC2 has been proposed as a tumor suppressor in breast cancer (Liu et al. 2020), but has not been explicitly linked to estrogen signaling. Similarly, GLIS3 is implicated in breast cancer, but not ER+ breast cancer (Rami et al. 2016). KLF5 has been shown to suppress ER in breast cancer (Guo et al. 2010), but has not been studied in endometrial cancer. The KLF9 and PAX8 motifs were found to be predictive of ER binding in Ishikawa cells only, and both have been shown to be implicated in endometrial cancer but have not been connected to ER activity specifically. KLF9 has been shown to alter cell morphology, proliferation, and differentiation in an ER- endometrial cancer cell line (Simmen et al. 2008), and high PAX8 protein levels are associated with poorer outcomes in endometrial cancer patients (Mhawech-Fauceglia et al. 2012).

Predicting regulatory EREs proved to be more challenging than predicting bound EREs. Fewer factors were found to be important, and the predictive models exhibited lower accuracy than those created to predict bound EREs. Despite the lower predictive ability, some strong patterns emerged. While transcription factor occupancy, ERE strength, and chromatin accessibility were the top predictors of ERE binding, chromatin looping and transcription were more predictive of regulatory EREs. Promoter-distal EREs that are engaged in transcription prior to ER binding are more likely to impact the expression of target genes. This is consistent with previous studies showing that transcription is a hallmark of active enhancer regions and that enhancer RNA production mirrors that of its target promoter (Field and Adelman 2020; Panigrahi and O’Malley 2021). Looping of an ERE both to a target gene promoter and other regulatory regions was also predictive of ERE regulatory activity. This is consistent with our previous study (Abewe et al. 2025), where we found that estrogen regulated genes with increased 3D genomic interactions after estrogen treatment had larger transcriptional responses, and most ER bound sites were anchors for chromatin loops. ER-mediated loops have also been identified in breast cancer and have been shown to change as the cancer progresses (Fullwood et al. 2009; Zhou et al. 2019; Van Den Brand et al. 2025). Overall, this leads to a model where bound EREs with high levels of transcription prior to estrogen treatment and many 3D genomic interactions are very likely to be regulatory; however, most regulatory EREs cannot be identified in this manner, suggesting that important features could be missing.

To experimentally evaluate the factors important for ERE activity, we used CRISPR KO to test the importance of thousands of epigenetic factors. We focused on Ishikawa cells because similar CRISPR-based techniques have been used to study ER control in breast cancer, and their results corroborate our findings (Xiao et al. 2018; Samuels et al. 2025). We encountered many histone modifying enzymes, which supported our computational findings. *ASH1L* encodes an H3K36me3 methyltransferase, which is related to transcription (An et al. 2011), *KAT6A* and *BRPF1* encode members of a histone acetylation complex associated with gene activation (Huang et al. 2016), and *KMT2A* encodes an H3K4me3 methyltransferase (Nakamura et al. 2002), a histone modification confirmed important by Boruta in both cell lines for predicting both bound and regulatory EREs. *BRPF1* has also been identified as a key component of estrogen signaling in ER+ breast cancer; silencing of *BRPF1* inhibits ER transactivation capability, and causes cell proliferation arrest and activation of apoptotic mechanisms (Salvati et al. 2024). *CITED2* has been identified as a transcriptional coactivator that modulates ER in breast cancer (Lau et al. 2013), indicating that some transcriptional coactivators are shared between breast and endometrial cancer. *EZH2* and *SUZ12* were both identified as ER enabling, which is unexpected as they are members of PRC2, a transcriptional repressive complex. It is possible that PRC2 represses genes that repress ER, making it an indirect supporter of ER activity; however, the role of polycomb proteins in cancer is controversial. *EZH2* and other members of PRC2 have been implicated as tumor suppressors in multiple hematopoietic cancers (Pasini and Di Croce 2016), but they have also shown oncogenic potential in multiple murine leukemia models and are being investigated as potential therapeutic targets (Scelfo et al. 2015). We identified *HDAC1* as an ER enabling gene, even though it is usually considered a corepressor (Hui Ng and Bird 2000). However, it has been positively associated with transcriptional activation caused by glucocorticoid receptor (Qiu et al. 2006), another steroid hormone receptor, so it could be associated with ER in a similar way and promote ER activity in endometrial cancer cells.

Previous work identified the SOX motif as uniquely enriched at ER binding sites in endometrial cancer cells (Rodriguez et al. 2020). Although there are many SOX family members, we chose SOX17 for further investigation as it is the highest expressed SOX factor in endometrial cancer cell lines, it is essential for murine endometrial gland development, it is involved in key aspects of endometrial biology such as cycling and implantation (Guimaraes-Young et al. 2015; Wang et al. 2018; Kinnear et al. 2019), and it has been identified as a potential biomarker for endometrial and ovarian cancers (Shaker et al. 2023). Because of this, we performed ChIP-seq on SOX17 and were able to establish that SOX17 binding is predictive of bound EREs in Ishikawa cells. These results were supported by our identification of *SOX17* as an ER enabling gene in the Ishikawa CRISPR KO screen.

By performing *SOX17* KO, we were able to validate that SOX17 is important for ER activity in endometrial cancer cells. *SOX17* KO caused significant reductions in ER reporter activity, ER ChIP-seq signal, and gene expression responses to estrogen. SOX17 likely regulates ER activity in two ways. SOX17 plays a role in activating *ESR1* transcription and driving higher protein levels by binding near the gene. SOX17 also binds to thousands of ER binding sites across the genome in the absence of estrogen, potentially aiding in ER genomic binding. In endometrial cancer patient samples, *SOX17* mRNA expression is positively correlated with *ESR1* mRNA expression, and *SOX17* is overexpressed in tumor samples compared to normal samples, partially through genomic amplification in 4.7% of tumors. *SOX17* is altered in ∼11-15% of endometrial tumors, although these alterations do not seem to have an effect on overall expression levels, and the tumors included in these studies are not all ER-positive (Levine 2013; Walker et al. 2017). SOX17 has also been identified as a tumor suppressor in endometrial cancer (Zhang et al. 2016; Walker et al. 2017), indicating a complex role for SOX17 in endometrial cancer that may go beyond its effects on estrogen signaling.

While this study has uncovered interesting features of ER genomic binding and gene regulation, there are limitations that should be mentioned. For the machine learning analysis, we considered only one ERE at a time, despite our previous work showing that EREs often work together to regulate gene expression (Ginley-Hidinger et al. 2019; Carleton et al. 2020; Ginley-Hidinger et al. 2024). We focused solely on full EREs even though half sites have been shown to be present at about 38% of ER binding sites in T-47D and Ishikawa cells (Gertz et al. 2013) and these sites can functionally impact gene regulation (Carleton et al. 2020). ERE half sites are six base pairs long and would be expected to occur more than 700,000 times in the human genome by random chance, making it computationally infeasible to evaluate these sites in the same way that we evaluated the full ERE sites. We classified bound EREs within 100kb of an estrogen responsive gene’s TSS as regulatory based on our previous analysis; however, we know that several of these sites do not impact gene expression (Carleton et al. 2017; Abewe et al. 2025). Functional studies of EREs in their genomic context would be the ideal method of identifying regulatory EREs; however, this has not been done at scale. Indicative of this issue, similar features were associated with regulatory EREs for both upregulated and downregulated genes, which is likely due to most regulatory EREs associated with downregulated genes being associated with upregulated genes as well. In addition, we were unable to achieve high sensitivity when predicting regulatory EREs. The data used in our machine learning analysis to compare Ishikawa and T-47D cells came from multiple sources. It is possible that differences discovered between the cell lines were due to technical artifacts as opposed to underlying biological differences. For example, the ATAC-seq data came from two different studies, and our analyses showed that ATAC-seq was a better predictor of bound EREs in Ishikawa cells than it was in T-47D cells. This is likely a technical artifact, as previous literature has established that the chromatin environment is important for ER binding in both cell lines (Gertz et al. 2013). The results from the CRISPR KO screen provided valuable information to support our computational results, but it also has limitations. With the reporter readout, we are unable to disentangle factors that affect ER binding and factors that impact ER’s regulatory activity. GFP reporter expression levels were already high in response to estrogen, making it difficult to identify genes that decrease ER activity. While the CRISPR screen and the machine learning analyses identified interesting factors, further studies are needed to determine the underlying mechanisms and functional importance of each factor. The hope is that our findings motivate future studies into important gene regulation factors in estrogen driven endometrial cancer.

## Methods

### Estrogen response element identification and classification

To identify all EREs in the human female genome, the program Patser-v3e (Hertz and Stormo 1999) was used to scan the hg19 genome to find and score sequences that matched closely to the canonical ERE motif, as defined by a position-specific scoring matrix (PSSM) created from the binding sites of ER in multiple estrogen responsive cell lines (Gertz et al. 2013). The hg19 build of the human genome was used to align with previously published datasets in our analyses. We do not believe that realigning our data to the GRCh38 genome build would affect the results of this work, as we focused our analyses on uniquely alignable regions of the genome. Patser was run with the following flags "-c -u2 -v". Regions above the numerically calculated cutoff were kept for downstream analysis as EREs, and the score was kept as a representation of the log of the relative affinity that region has for binding ER. The default cutoff score is the score at which the probability of observing a score greater than or equal to the cutoff is equal to the adjusted information content (Guhathakurta 2006). The adjusted information content is calculated based on the divergence of frequency of each nucleotide within a sequence from the a priori frequency of each nucleotide (each nucleotide occurring 25% of the time) and adjusted based on the length of the consensus sequence (the ERE) and the size of the sequences being scanned (the human genome) (Hertz and Stormo 1999). Since both cell lines are female, any EREs found on the Y Chromosome were removed from this list. The list of EREs was intersected, using BEDtools intersect (Quinlan and Hall 2010), with published ER ChIP-seq data taken from either T-47D cells or Ishikawa cells (Gertz et al. 2013). Any ERE that intersected with an ER ChIP-seq peak was labeled as “bound”; otherwise, it was labeled as “unbound”. Then, significant genes from published bulk RNA-seq data (Gertz et al. 2013) and single-cell RNA-seq data (Ginley-Hidinger et al. 2024) were combined to create a list of estrogen responsive genes for each cell line. For the bulk RNA-seq with an 8-hour estrogen treatment, any gene with an adjusted *P*-value of 0.05 or lower was included as an estrogen responsive gene. For the single-cell RNA-seq data, any gene that had an adjusted *P*-value of 0.05 or lower at any time point (2-, 4-, or 8-hour) of estrogen treatment compared to the 0-hour timepoint was included as an estrogen responsive gene. The final list of estrogen responsive genes was the union of genes from the bulk RNA-seq and the single-cell RNA-seq. A bound ERE within 100kb of the TSS of an estrogen responsive gene was labeled as “regulatory”. A bound ERE within 5kb of the TSS of an estrogen responsive gene also received the designation of promoter-proximal, with all other regulatory EREs receiving the designation of promoter-distal.

### Comparison of bound EREs in cell line models to bound EREs in patient-derived models

All significant EREs were intersected with ER ChIP-seq data from multiple breast and endometrial cancer patient samples (Rodriguez et al. 2020) using BEDtools intersect (Quinlan and Hall 2010). Then, bound EREs from each sample were intersected with the bound EREs in Ishikawa and T-47D cells. A Wilcoxon sign-ranked test was performed to compare the overlaps between each patient sample and cell line to determine if Ishikawa bound EREs and T-47D bound EREs were more similar to endometrial and breast cancer patient samples, respectively.

### Creation of experimental-focused dataset and motif-focused dataset for feature selection analysis

For the initial data frames used, the relative affinity score given by Patser, ChIP-seq of multiple transcription factors and histone modifications, ATAC-seq, DNase-seq, and PRO-seq were compiled. For each ERE, the normalized reads per million of each ChIP-seq, ATAC-seq, DNase-seq, and PRO-seq dataset at the location of the ERE were calculated, and then each dataset was converted to *Z*-scores. See Supplemental Table 1 for information about all datasets. All data were from T-47D and Ishikawa cells grown in the absence of estrogen. HiChIP data was used to determine which bound EREs had a 3D genomic interaction with the 5kb window around the TSS of an estrogen responsive gene, and this data factor was labeled “Target Gene 3D Loop.” HiChIP was also used to determine the number of 3D genomic interactions in which each ERE (bound or unbound) was involved. To prevent redundancy with the Target Gene 3D Loop data, we only counted interactions that did not involve the TSS of an estrogen responsive gene. This factor was called “Non-Target Gene Loops.”

For the motif analysis, the comprehensive hg19 BED file from Homer that lists the locations of the cognate DNA binding motifs for 450 DNA-binding proteins was used (Heinz et al. 2010). BEDtools intersect (Quinlan and Hall 2010) was used to intersect this file with a BED file of the EREs in the human female genome with a 100bp window centered on each ERE. A binary classifier was then created for each motif. An ERE either had the motif in its 100bp window or it did not. Spearman correlation between each motif was calculated using the R command cor from the stats package (R Core Team 2016). Using the correlation values, clustering using hclust from the R stats package, with the average method, was performed, and the cluster tree was cut off at 0.50, reducing the list of motifs down to 270 motif clusters (Table S3). The binary classifier was updated to reflect whether an ERE 100 bp window had any of the motifs within each cluster. Motif clusters that contained a motif that matched at least an ERE half-site were not displayed or considered for further analysis.

### Feature selection analysis using Boruta

The default settings were used for all Boruta runs, using the function Boruta from the R package Boruta (Kursa and Rudnicki 2010), except that the number of threads used was 20. For all initial runs, the entire dataset was used, meaning all EREs. In addition, Boruta was performed using down-sampling for predicting ER binding. All bound EREs were used with an equal number of randomly selected unbound EREs to account for the much larger number of unbound EREs. This was repeated 100 times, and the average importance scores and the percentage of runs where the factor was confirmed important by Boruta were calculated. Down-sampling was not used when running Boruta to evaluate predicting effects on expression, because the number of regulatory and nonregulatory EREs was similar.

### Creation of receiver operating characteristic curves and logistic regression models

The command roc, from the package pROC (Robin et al. 2011), was used with default parameters to create the receiver operating characteristic curves (ROC) and calculate the area under the curve (AUC). The R package caret (Kuhn et al. 2020) was used to perform logistic regression. To set up training and testing groups, the command createDataPartition was used, with the training groups set as 75% of the data and testing groups as 25%, repeated for 100 iterations. To set parameters for the training, the command trainControl was used, using the index from createDataPartition with the following parameters: the method was set to cross validation (“cv”), class probabilities were set to be calculated instead of just the final class (classProbs=T), the summary function was set to “twoClassSummary”, meaning the ROC AUC, the sensitivity, and the specificity were designated as the metrics for assessing the performance of the model, and down-sampling was designated to account for the disparities in the dataset so that the training model used equal amounts of each type of ERE. The command train was used with the above training parameters, with logistic regression implemented using the options: method = “glm” and family = “binomial”. The models returned the average and standard deviation of ROC AUC, sensitivity, and specificity from multiple iterations. This approach was used for creating models for both bound vs unbound EREs and regulatory vs. nonregulatory EREs, for all cell lines, with different variables depending on the application. Bound EREs were set to 1, and unbound EREs were set to 0; similarly, regulatory EREs were set to 1, and nonregulatory EREs were set to 0, with the logistic regression models using the default cutoff value of 0.5.

### Cell culture

Ishikawa, T-47D, and HCI-EC-23 cells were grown in full media: RPMI 1640 medium (Thermo Fisher Scientific, 11875135) with 10% fetal bovine serum (Thermo Fisher Scientific, 26140079) and 1% penicillin–streptomycin (Thermo Fisher Scientific, 15070063). Hormone depleted media was also used in preparation for hormone treatment and consisted of phenol red-free RPMI (Life Technologies, 11835055) with 10% charcoal-dextran-stripped fetal bovine serum (Sigma-Aldrich, F6765-100ml) and 1% penicillin–streptomycin (Thermo Fisher Scientific, 15070063). Cells were incubated at 37°C with 5% CO_2_.

### ChIP-seq

Cells were grown in hormone depleted media for 5 days prior to either a 1 hour DMSO (Fisher Scientific, BP231) (vehicle) treatment (SOX17), a 1 hour 10 nM 17β-estradiol (E2) (Sigma Aldrich, E2758-1G) treatment (ER), or an 8-hour DMSO treatment (H3K4me1), and were harvested for ChIP-seq (Johnson et al. 2007) as described previously (Reddy et al. 2009). Cells were fixed using 1% formaldehyde (Sigma Aldrich, 252549) for 10 minutes at room temperature and then quenched using glycine (Fisher Scientific, BP381), added to achieve a final concentration of 125 mM. Fixed cells were washed with cold PBS and harvested via scraping in PBS. Cells were pelleted at 500 RCF for 5 minutes, and PBS was removed. The cell pellet was resuspended in 1 mL of Farnham lysis buffer (5 mM PIPES pH 8 (Fisher Scientific, 50-155-760), 85 mM KCl (Sigma-Aldrich, P9541), and 0.5% NP-40 (VWR, M158-100ML)) supplemented with protease inhibitors (Thermo Scientific, A32965) to prepare crude nuclei. Nuclei were pelleted at 500 RCF for 5 minutes and Farnham lysis buffer was removed. Nuclei were resuspended in RIPA lysis buffer (1× PBS (Thermo Scientific, AM9624), 1% NP-40 (VWR, M158-100ML), 0.5% sodium deoxycholate (Bioworld, 40430018-3), and 0.1% SDS (Thermo Scientific, AM9820)) supplemented with protease inhibitors (Thermo Scientific, A32965) and sonicated at 4°C for 8 cycles (30 seconds on, 30 seconds off) at 40% amplitude using a EpiShear probe sonicator (Active Motif). Chromatin immunoprecipitation was performed using 5 μg of antibody. Antibodies used for this study were SOX17 (R&D Systems, AF1924), ER (Abcam, AB3575), and H3K4me1 (Cell Signaling, 5326S). ChIP-seq libraries were sequenced on an Illumina NovaSeq 6000. Reads were aligned to the hg19 build of the human genome using Bowtie (Langmead et al. 2009): -m1 -t -best -q -S -l 32 -e 80 -n 2. MACS2 (Zhang et al. 2008) was used to call peaks with a *P*-value cutoff of 1×10^-10^ and mfold parameter between 15 and 100.

### ATAC-seq

HCI-EC-23 cells were grown in hormone-depleted media for 5 days prior to a 1-hour DMSO treatment and harvest for ATAC-seq (Buenrostro et al. 2013). Cell pellets were treated and lysed for 5 minutes as described previously (Lewis et al. 2025). The transposition reaction, DNA purification, and amplification were also performed as described previously (Lewis et al. 2025). Libraries were sequenced on an Illumina NovaSeq X at a depth of ∼50 million reads/sample and reads were trimmed to 50 bp. Bowtie (Langmead et al. 2009) was then used to align the trimmed reads to the hg19 build of the human genome, using the following parameters: -m 1 -t --best -q -S −l 32 -e 80 -n 2. BAM files were created from SAM files and sorted using SAMtools (Li et al. 2009). Regions of accessible chromatin, termed peaks, were called using a cutoff of <0.05 for adjusted *P*-value and an mfold parameter between 15 and 100 using MACS2 (Zhang et al. 2008). Reads that aligned to the peaks were quantified across all samples using featureCounts (Liao et al. 2014).

### Generation of 3×ERE-driven GFP lentiviral plasmid

The pLU-CMV-GFP-pBLAST plasmid, a gift from Meenhard Herlyn (Kerafast, EW1034) (Roesch et al. 2010), was used as the destination lentiviral plasmid. pLU-CMV-GFP-pBLAST’s CMV promoter was removed by restriction digest using ClaI (NEB, R0197S) and XbaI (NEB, R0145S), with 3’-phosphate groups removed using shrimp alkaline phosphatase (NEB, M0371S). The backbone was size selected using gel extraction (Qiagen, 28704). A 3×ERE sequence was PCR amplified from a donor plasmid, 3×ERE TATA Luc, a gift from Donald McDonnell (Addgene, 11354) (Hall and McDonnell 1999), using primers that added directional ClaI and XbaI restriction sites (Table S6). PCR products were digested using ClaI and XbaI, purified using DNA Clean and Concentrator (Zymo Research, D4014), and ligated into the pLU-CMV-GFP_pBLAST backbone. Ligation products were transformed into STBL3 chemically competent cells (Thermo Scientific, C737303) and selected on ampicillin (100 μg/mL) LB agar plates. Correct insertion of the 3×ERE sequence was determined using Sanger sequencing (Genewiz), and the resultant plasmid was termed pLU-3×ERE-GFP-pBLAST. Lentiviral particles were generated by co-transfection of pLU-3×ERE-GFP-pBLAST with helper plasmids pCMV-VSV-G, a gift from Bob Weinberg (Addgene, 8454) (Stewart et al. 2003), and psPAX2, a gift from Didier Trono (Addgene, 12260) into HEK293T cells.

### Ishikawa GFP reporter cell line creation

Ishikawa cells were transduced with 5 μg/mL polybrene (Santa Cruz Biotechnology, sc-134220) and the lentiviral particles containing the 3×ERE-driven GFP reporter at MOIs of 10-200. Cells were treated with 30 μg/mL blasticidin (Thermo Fisher Scientific, A1113903) initially to select for successfully transduced cells and then pooled. After 96 hours, the blasticidin dose was reduced to 10 μg/mL for maintenance, and 10 nM E2 was added to activate GFP. Cells were incubated in full media for 72 hours and then sorted for positive GFP expression using flow cytometry. GFP positive cells were plated at 1000 cells per 10cm plate to allow single cells to form distinct colonies for selection. Colonies were grown up to ∼256 cells on media without blasticidin or E2, then selected and moved to individual wells of a 96 well plate. 10 μg/mL blasticidin was added to full media 48 hours after colonies were plated in 96 well plates and included in media for all further culturing of the cells.

To test GFP responsiveness, colonies were moved to hormone depleted media for 72 hours, then treated with 10 nM E2 or DMSO (vehicle) for an additional 72 hours. Then the GFP levels of the cells were measured using flow cytometry, and colonies with the largest difference in GFP levels between E2 and DMSO conditions were chosen to use for further experimentation. To increase the GFP fold change for the chosen colony, cells were treated with 100 μg/mL blasticidin for two weeks, and then retested for GFP responsiveness before being used in the CRISPR screen.

### CRISPR epigenetic and transcription factor KO screen

The Human Epigenetic Knockout plasmid library (HEKL), which was a gift from Kivanc Birsoy (Addgene 162256) (Williams et al. 2020), was co-transfected with helper plasmids pCMV-VSV-G, a gift from Bob Weinberg (Addgene, 8454) (Stewart et al. 2003), and psPAX2, a gift from Didier Trono (Addgene, 12260), into HEK293T cells to generate lentiviral particles. Ishikawa cells with the 3×ERE GFP reporter were grown to confluency in 3 15cm plates. One plate was used to count cells, and the other two plates were transduced at an effective MOI of 0.3 with the HEKL lentiviral particles and 5 μg/mL polybrene in full media. Puromycin was used at 1μg/mL in full media to select for successfully transduced cells 24 hours after transduction. Post puromycin selection, the cells were grown up to 10 million cells in full media with 10 μg/mL blasticidin, 2.5 million cells were frozen down for genomic DNA extraction and sequencing, and 2.5 million cells were moved to hormone depleted media with 10 μg/mL blasticidin for 96 hours. After 96 hours in hormone depleted media, 2.5 million cells were frozen for genomic DNA extraction and sequencing, and the remaining cells were treated with 10 nM E2 for 72 hours. Cells were then sorted using flow cytometry into four bins: negative GFP, low GFP, medium GFP, and high GFP. Genomic DNA was extracted from the cells in all four bins along with the cells from the post transduction pellet, and the post-hormone depleted media pellet using the Quick-DNA Miniprep Kit (Zymo, D3025). Libraries were constructed for sequencing the CRISPR sgRNA region using a two-step PCR process. Genomic DNA (100 to 5000 ng) was PCR amplified for 14 cycles using custom CRISPR PCR primers with NEBNext Ultra II Q5 Master Mix (NEB, M0544) (Table S6). This was performed as 8 replicate reactions with the amplicons being pooled before the next step. Dual unique indexing adapters (Illumina, 20027213) (Table S6) were added to the CRISPR amplicons during a second PCR with 6 cycles and performed as 3 replicate reactions. Libraries were pooled and sequenced on an Illumina NovaSeq X instrument as 150×150 cycle paired-end sequence runs. Two replicates, starting with transduction of the HEKL lentiviral particles, were performed.

### Post-screen enrichment analysis using MAGeCK

The forward read FASTQ files, along with a guide RNA file, were analyzed using the MAGeCK program (Li et al. 2014; Williams et al. 2020). MAGeCK count was used to compile counts of each gRNA in each sample. This command allowed for combining two replicates to generate a single count file. MAGeCK test was used to test for enrichment. The first run designated the negative GFP bins as the control with the other three as the samples and the second run designated the negative, low, and medium GFP bins as the controls with the high GFP bin as the sample. After testing, the gene outputs were filtered by the number of gRNAs used in analysis, designated by MAGeCK as “good gRNAs.” Genes included for future analysis had to have more than half of their gRNAs included in the test analysis performed by MAGeCK. After filtering, the command p.adjust from the R stats package was used to adjust the *P*-values, using the “fdr” method. The negative significant genes from the negative GFP control analysis were labeled as ER enabling genes, and the positive significant genes from the negative, low, and medium GFP control analysis were labeled as ER blocking genes.

### Ishikawa Cas9 cell line creation

The lentiCas9-Blast plasmid, a gift from Feng Zhang (Addgene, 52962) (Sanjana et al. 2014) was co-transfected with helper plasmids pCMV-VSV-G, a gift from Bob Weinberg (Addgene, 8454) (Stewart et al. 2003), and psPAX2, a gift from Didier Trono (Addgene, 12260), into HEK293T cells to generate lentiviral particles. Ishikawa cells were then transduced with 5 μg/mL polybrene and the lentiviral particles containing the 3×ERE-driven GFP reporter at MOIs of 0.5-10. Cells were treated with 30 μg/mL blasticidin initially to select for successfully transduced cells; after selection, the blasticidin dose was reduced to 10 μg/mL for establishment prior to cryo-preservation. Cas9 presence was verified using Western blot and an MOI of 5 was chosen.

### Creation of ER and SOX17 targeting lentiviral plasmids

Guide RNAs (gRNAs) were designed to target *ESR1* and *SOX17* and ordered as single-stranded DNA primers from IDT (Table S6). For a control, we used the control sequence used previously (Sanjana et al. 2014). The LentiGuide-Puro plasmid, a gift from Feng Zhang (Addgene, 52963) (Sanjana et al. 2014), was used as the destination lentiviral plasmid. The plasmid was restriction digested using FastDigest Esp31 (Thermo Fisher Scientific, FD0454) with 3’-phosphate groups removed using shrimp alkaline phosphatase (NEB, M0371S). The backbone was size selected using 0.8% gel extraction. The gRNA primers were annealed in 10× T4 Ligation Buffer (NEB, B0202S) and phosphorylated using T4 Polynucleotide Kinase (NEB, M0201S) before being ligated to the destination plasmid using the Quick Ligation Kit (NEB, M2200S). Ligation products were transformed into STBL3 chemically competent cells (Thermo Scientific, C737303) and selected on ampicillin (100 μg/mL) LB agar plates. Correct insertion of gRNAs was determined using full plasmid sequencing (Plasmidsaurus). Lentiviral particles were generated by co-transfection of the LentiGuide Puro plasmids (with the gRNAs inserted) with helper plasmids pCMV-VSV-G, a gift from Bob Weinberg (Addgene, 8454) (Stewart et al. 2003), and psPAX2, a gift from Didier Trono (Addgene, 12260) into HEK293T cells.

### Creation of *ESR1* KO, *SOX17* KO, and control gRNA lines

Ishikawa Cas9 expressing cells, described above, were transduced with 5 μg/mL polybrene and the lentiviral particles containing the LentiGuide Puro plasmids at MOIs of 10-200. Cells were treated and maintained with 1μg/m puromycin in full media until validation. Western blotting was used to confirm the absence of ER and SOX17 in their respective KO lines after puromycin selection. After Western blot confirmation, cells were plated at 1000 cells per 10 cm plate to allow single cells to form distinct colonies for selection. Colonies were grown up to ∼256 cells on full media, then selected and moved to individual wells of a 96 well plate before further passaging. Individual clones of the KO lines were evaluated for ER and SOX17 absence using Western blotting.

### Western blotting

Western blotting was performed on Ishikawa control cells, Ishikawa Cas9 cells, *ESR1* KO cells, and *SOX17* KO cells as described previously (Rush et al. 2022) using the following primary antibodies: mouse anti-β-Actin (Santa Cruz Biotechnology, sc-47778), rabbit anti-Cas9 (Diagenode. C15310258), rabbit anti-ERα (Thermo Fisher Scientific, MA5-14501), goat anti-SOX17 (R&D Systems, AF1924). The following secondary antibody conjugates were used for detection: IRDye 680RD donkey anti-Mouse (Li-COR, 926-68072), IRDye 800CW donkey anti-rabbit (Li-COR, 926-32213), and IRDye 800CW donkey anti-Goat (Li-COR, 926-32214). Transfer onto nitrocellulose membranes (Thermo Scientific, IB33002X3) was performed using the iBlot 3 dry blotting system (Thermo Scientific, IB31001). Fluorescence signal from the antibodies were then detected using the Li-COR Odyssey Clx imaging system with the 800CW and 680RD channels. Western blot images were analyzed using Image Studio software (Li-COR).

### ERE luciferase reporter assay

Ishikawa *SOX17* KO, *ESR1* KO, and control cells were grown in hormone depleted media for 3 days prior to plating at 15,000 cells per well in a 96-well plate in hormone depleted media. Then, the ERE luciferase reporter assay was performed as described previously using 10 nM E2 treatments (Rush et al. 2022).

### RNA-seq

Ishikawa *SOX17* KO and control cells were treated with 10 nM E2 or DMSO (vehicle) for 8 hours. Following treatment, cells were lysed with buffer RLT Plus (Qiagen) containing 1% beta-mercaptoethanol (Sigma-Aldrich). RNA was extracted and purified using a Quick RNA Mini Prep kit (Zymo Research). NEBNext Ultra II Directional RNA Library Prep kit with poly(A) mRNA isolation was used to construct RNA-seq libraries according to the manufacturer’s instructions (NEB). Sequencing reads were aligned to hg19 build of the human genome using HISAT2 (Kim et al. 2019). SAMtools (Li et al. 2009) was used to convert SAM files to BAM files. Genes were defined by the University of California Santa Cruz (UCSC) Known Genes (Kent et al. 2002) and reads that mapped to known genes were assigned with featureCounts (Liao et al. 2014). Read counts were normalized and analyzed for differential expression via DESeq2 (Love et al. 2014).

## Data access

All raw and processed sequencing data generated in this study have been submitted to the NCBI Gene Expression Omnibus (GEO; https://www.ncbi.nlm.nih.gov/geo/) under accession numbers GSE312322, GSE312323, GSE333956, GSE334097, and GSE334100.

## Acknowledgements

Funding for this work came from the National Institutes of Health (NIH)/National Cancer Institute (NCI) R21 CA270580 to J.G. and from the Huntsman Cancer Institute. Research reported in this publication utilized the High-Throughput Genomics Shared Resource at the University of Utah and was supported by NIH/National Cancer Institute (NCI) award P30 CA042014. The Flow Cytometry Shared Resource and the Drug Discovery Shared Resource at the University of Utah were also utilized during this study. We thank Matthew Ginley-Hidinger for his help with coding and figure making, Wilbert Zwart and Sebastian Gregoricchio for their helpful suggestions, Jacob Polaski for his guidance in designing the ATAC-seq protocol, and other Gertz lab members for their input on the study and manuscript.

## Author Contributions

Conceptualization was by A.R. and J.G. Methodology was by A.R., C.M.R., N.K., H.A., B.K.D., and J.G. Investigation was by A.R., N.K., C.M.R., H.A., K.S.O., and A.J.B. Formal analysis was by A.R., J.M.V., and J.G. Writing of the original draft was by A.R. and J.G. Writing, reviewing, and editing were by all authors. Supervision was by J.R. and J.G. Funding acquisition was by J.G.

