## Supplementary Figures and Tables 1 and 6 for "Identification of genomic features that uniquely impact estrogen receptor alpha binding and its effects on gene expression in endometrial cancer"

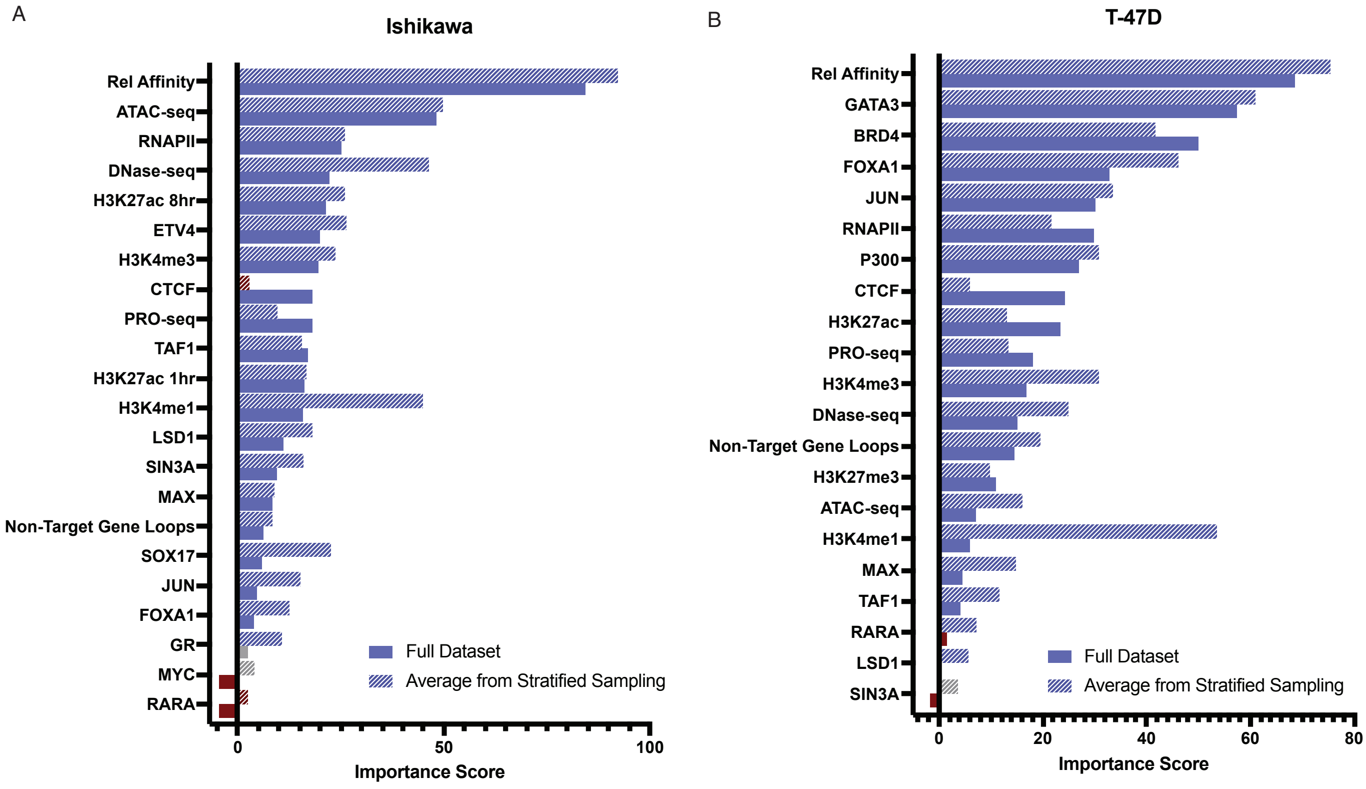


**Supplemental Figure 1: Down sampling produces similar Boruta results as the full dataset for ERE binding prediction.** Bar graph shows Boruta importance score from using full dataset (solid bars) compared to average importance score from down-sampled data (hashed bars) in A) Ishikawa and B) T-47D. The color of the bars indicates the decision; blue indicates confirmed in full data set Boruta run or confirmed in 80% or more of down-sampled Boruta runs, gray indicates tentative in full data set Boruta run or confirmed in between 50 and 80% of down-sampled Boruta runs, and red indicates rejected in full data set Boruta run or confirmed in less than 50% of down-sampled Boruta runs. This figure is related to Figure 2.

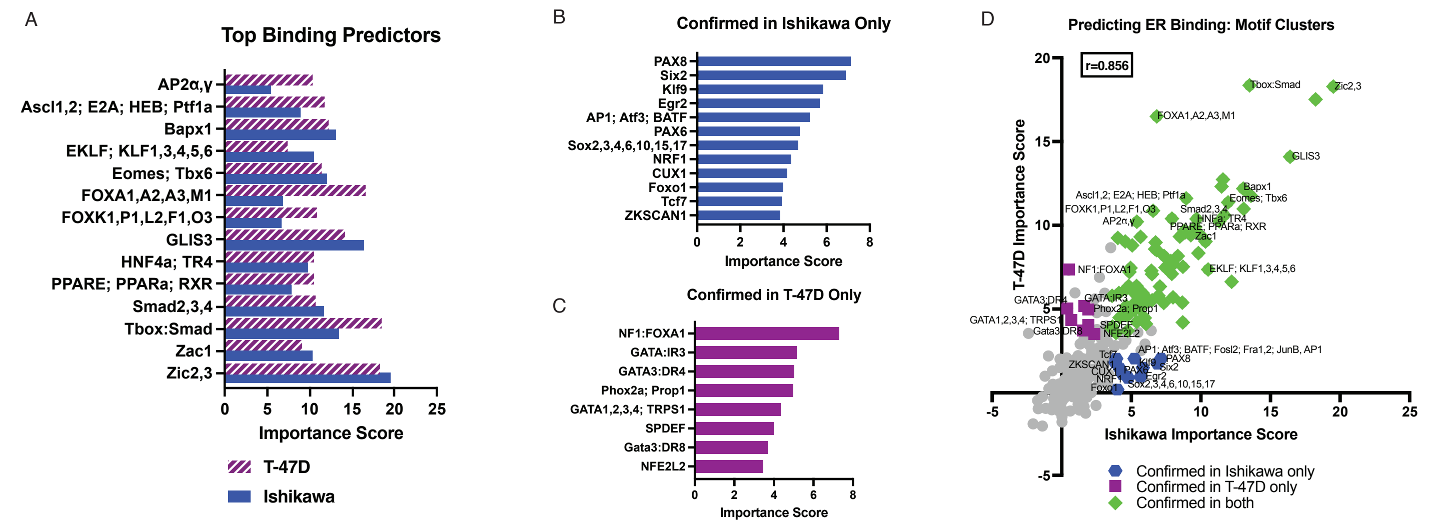


**Supplemental Figure 2: Motif analysis reveals that top motif predictors are shared between cell lines.** A) Bar chart shows the Boruta importance scores of the top scoring motif clusters for predicting bound EREs at an ERE in Ishikawa (blue) and T-47D (purple). Boruta importance scores are represented for motif clusters uniquely confirmed to be important in predicting bound EREs in B) Ishikawa and C) T-47D. D) Scatter plots show the comparison of Boruta importance scores between cell lines in predicting bound EREs from the motif cluster data set, with a Pearson correlation coefficient of 0.856. Green diamonds are clusters confirmed as important in predicting bound EREs by Boruta in both cell lines, blue hexagons are clusters confirmed only in Ishikawa, and purple squares are clusters confirmed only in T-47D. This figure is related to Figure 2.

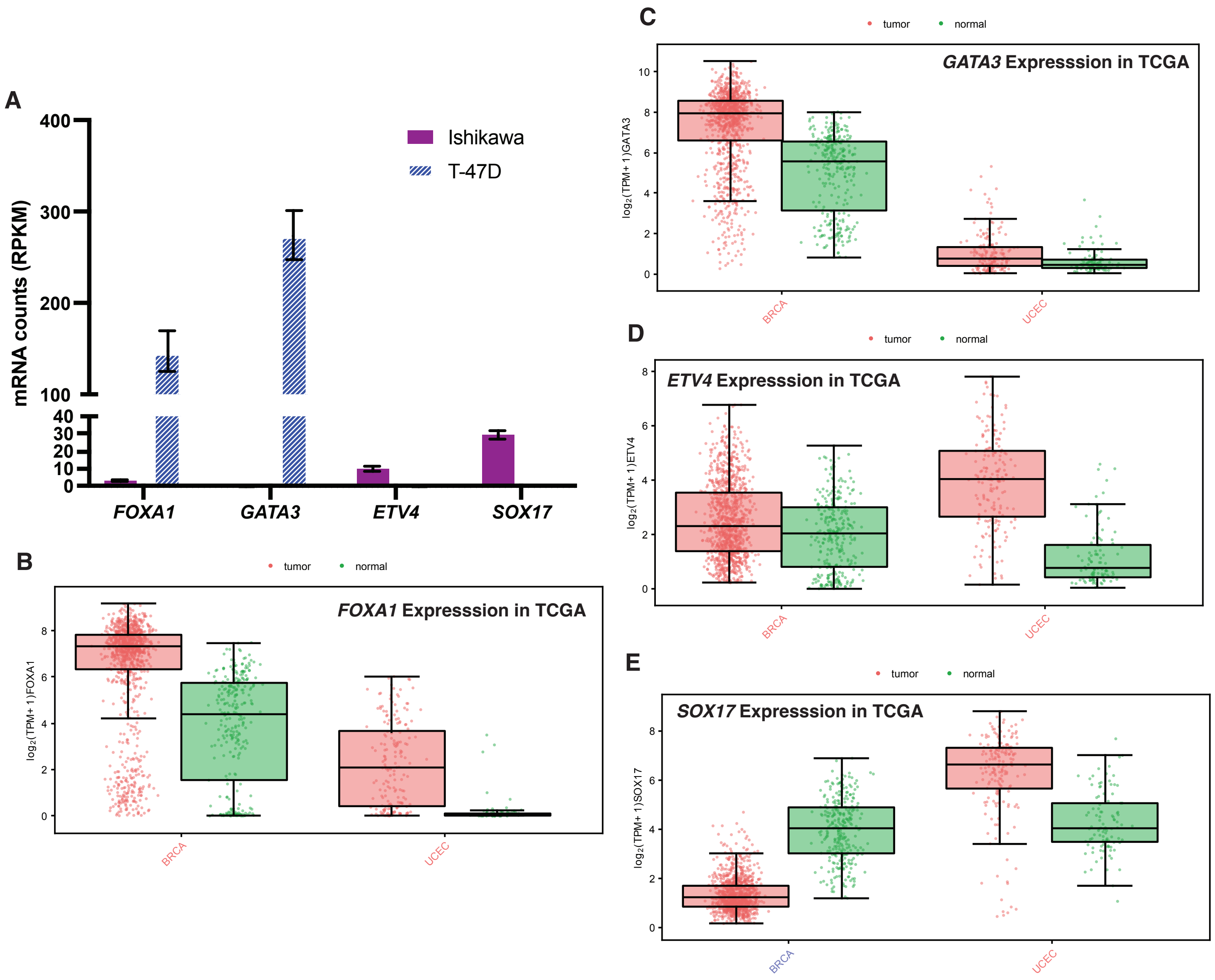


**Supplemental Figure 3: Cell type-specific transcription factors are expressed more highly in their corresponding tissue type.** A) Bar graph shows mRNA expression as reads per kilobase of transcript per million mapped reads for *FOXA1, GATA3, ETV4,* and *SOX17* in Ishikawa and T-47D cells. Boxplots show the expression of B) *FOXA1,* C) *GATA3*, D) *ETV4*, and E) *SOX17* in $\log_2$(TPM+1) in breast cancer samples (red, far left), breast normal tissue samples (green, second from left), endometrial cancer samples (red, second from right), and endometrial normal tissue samples (green, far right), created using GEPIA3 (Kang et al. 2025). This figure is related to Figure 2.

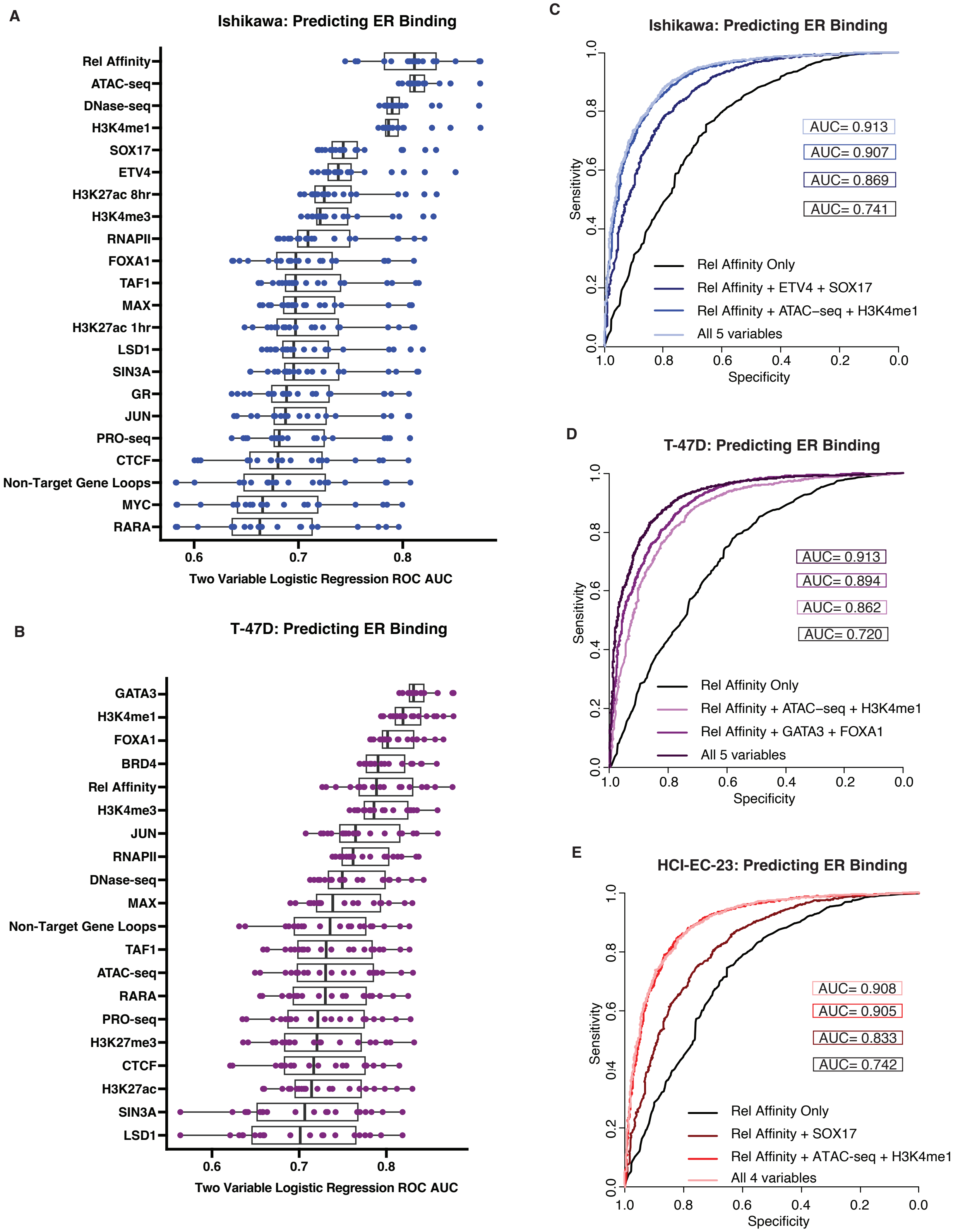


**Supplemental Figure 4: Multivariable logistic regression models predict bound EREs with high sensitivity and high specificity.** Ranked boxplots for each factor show the ROC AUC for all possible two variable logistic regression models predicting bound EREs in A) Ishikawa and B) T-47D that include that factor, i.e. the relative affinity boxplot is for all two variable logistic regression models containing relative affinity as one of the variables. Graphs show four different ROC curves, one using only relative affinity as a variable, one using relative affinity plus ATAC-seq and H3K4me1 ChIP-seq, one using relative affinity and cell line-specific transcription factor ChIP-seq datasets, and one using all variables from all other models for predicting bound EREs in C) Ishikawa, D) T-47D, and E) HCI-EC-23. AUC values are in boxes with colors corresponding to their ROCs.

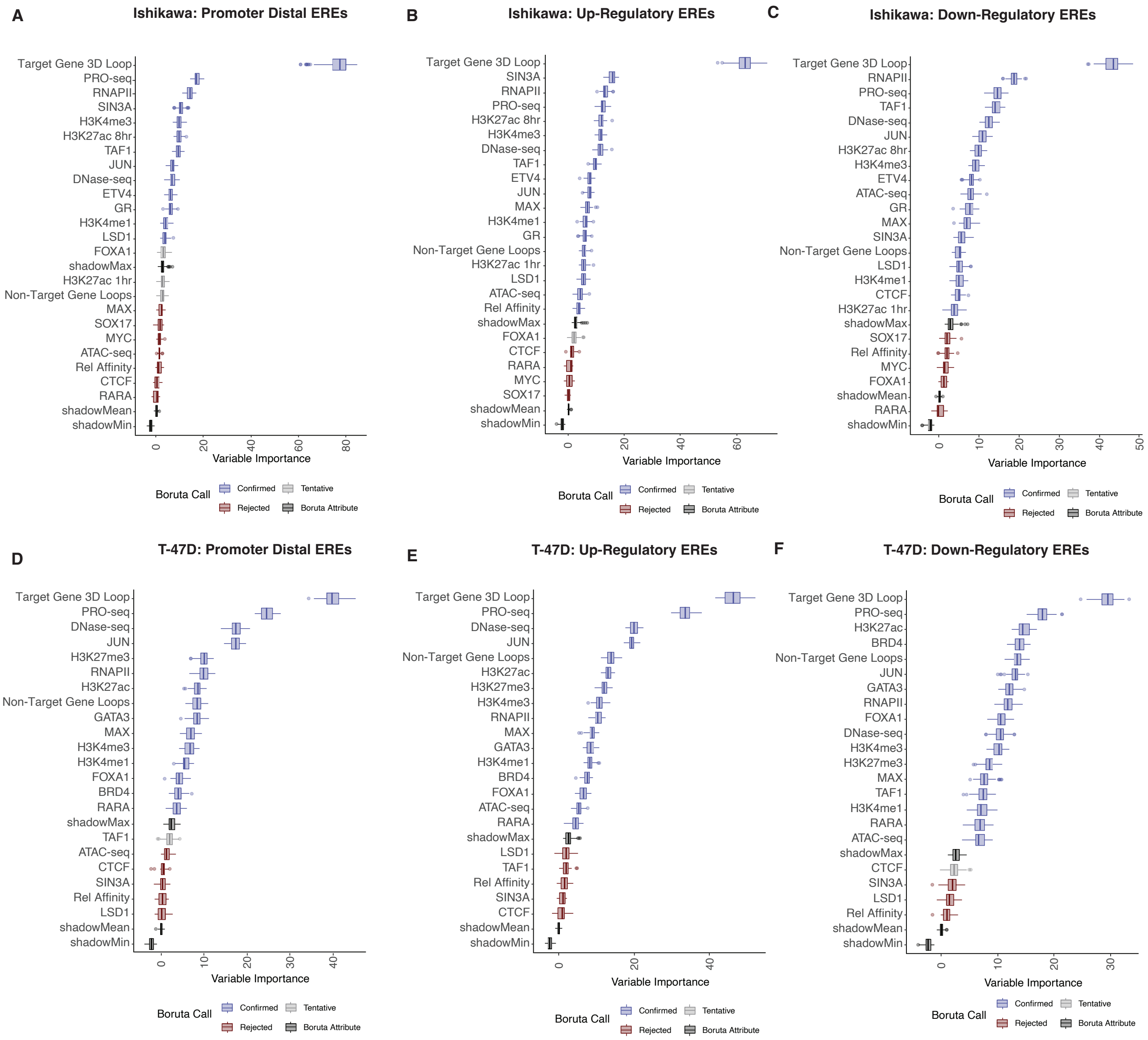


**Supplemental Figure 5: Gene distance and gene regulation direction minimally impact regulatory ERE prediction.** Boxplots show the ranked factor importance for predicting A) promoter-distal regulatory EREs versus nonregulatory EREs in Ishikawa, B) regulatory EREs associated with upregulated genes versus nonregulatory EREs in Ishikawa, C) regulatory EREs associated with downregulated genes versus nonregulatory EREs in Ishikawa, D) promoter-distal regulatory EREs versus nonregulatory EREs in T-47D, E) regulatory EREs associated with upregulated genes versus nonregulatory EREs in T-47D, and F) regulatory EREs associated with downregulated genes versus nonregulatory EREs in T-47D. Colors indicate Boruta determination of importance; blue is confirmed, light gray is tentative, red is rejected, and dark gray is a Boruta attribute. This figure is related to Figure 3.

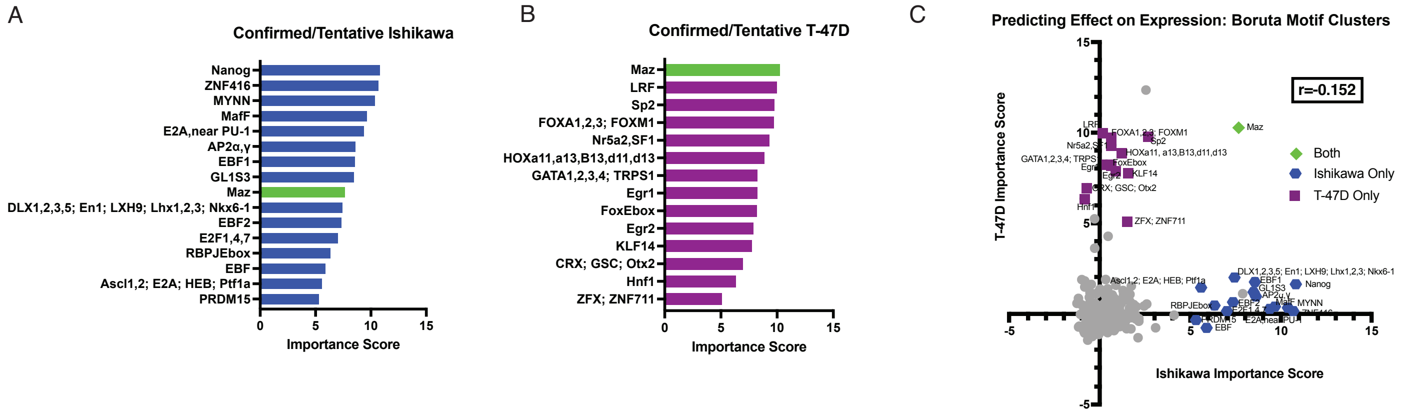


**Supplemental Figure 6: Regulatory EREs are poorly predicted using motif analysis.** Bar charts show motif clusters determined to be either confirmed or tentative by Boruta in being important for predicting regulatory EREs in A) Ishikawa or B) T-47D. Maz is the only motif confirmed important in both cell lines and is marked in green; all other motif clusters were uniquely determined to be confirmed or tentative in either Ishikawa (blue) or T-47D (purple). C) Scatter plots show the comparison of Boruta importance scores in predicting regulatory EREs from the motif cluster data set, with Pearson correlation coefficient of -0.152. The green diamond is the cluster confirmed as important in predicting regulatory EREs by Boruta in both cell lines, blue hexagons are clusters confirmed or designated tentative only in Ishikawa, and purple squares are clusters confirmed or designated tentative only in T-47D.  This figure is related to Figure 3.

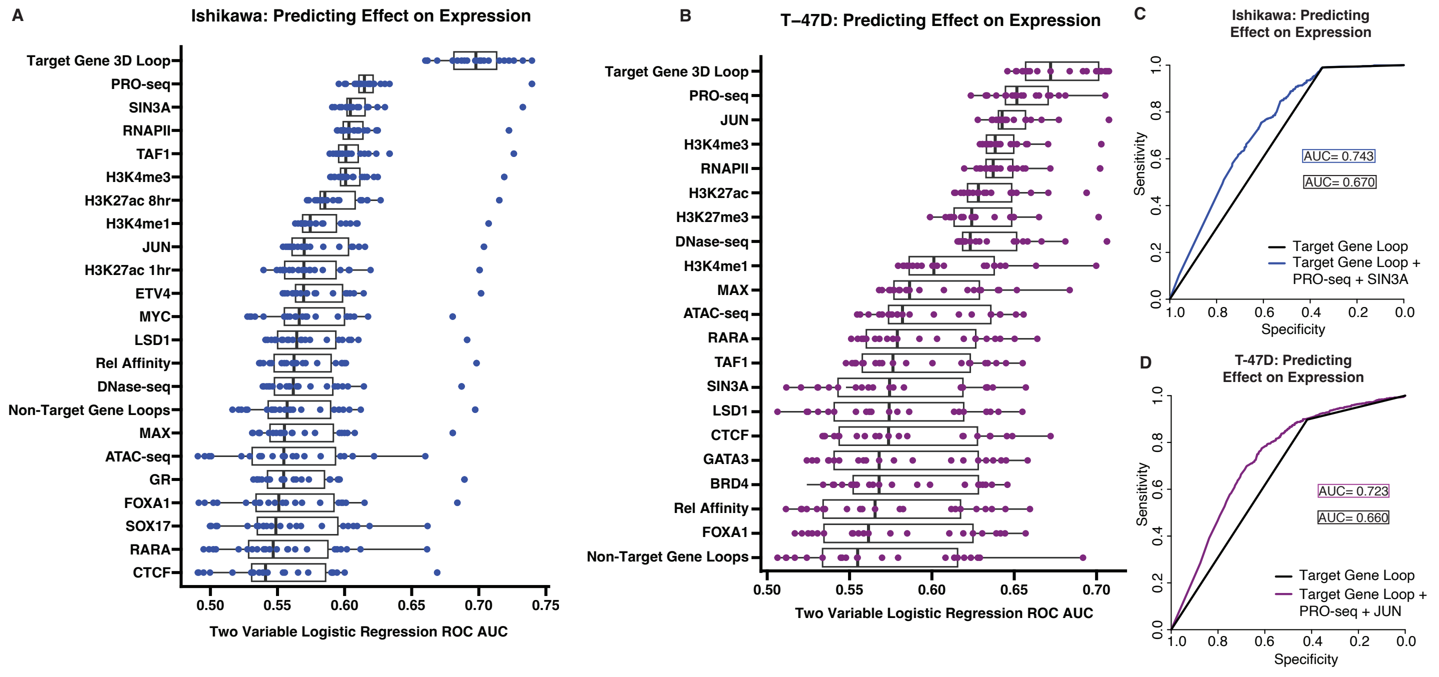


**Supplemental Figure 7: Multivariable logistic regression models predict regulatory EREs with high specificity, but low sensitivity**. Ranked boxplots for each factor show the ROC AUC for all possible two variable logistic regression models for predicting regulatory EREs that include that factor in A) Ishikawa and B) T-47D. Graphs show two different ROC curves, one using only Target Gene 3D Loop as a variable and one using Target Gene 3D Loop plus PRO-seq and a cell line-specific transcription factor ChIP-seq for predicting regulatory EREs in C) Ishikawa and D) T-47D. AUC values are in boxes with colors corresponding to their ROCs. This figure is related to Figure 3.

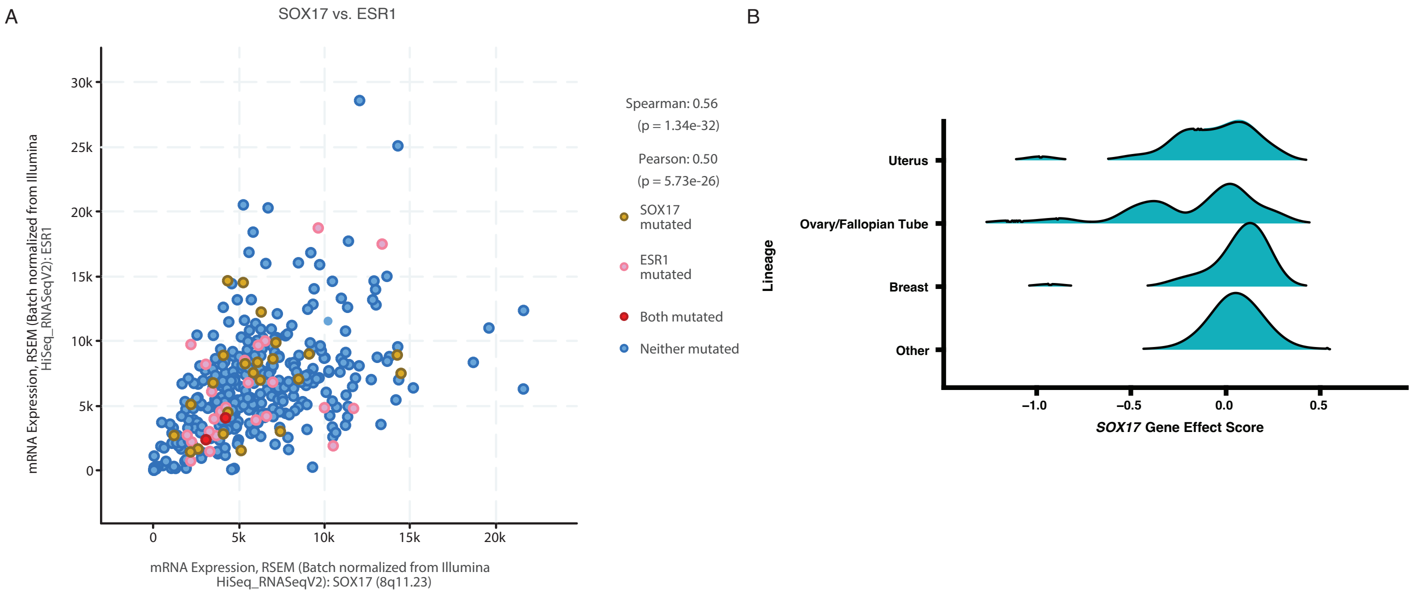


**Supplemental Figure 8: SOX17 expression in patient data samples from The Cancer Genome Atlas.** A) Scatter plot shows expression of *SOX17* versus the expression of *ESR1* in endometrial tumors using mRNA expression *Z*-scores relative to diploid samples, created using cBioPortal (Cerami et al. 2012; Gao et al. 2013; de Bruijn et al. 2023). B) Density plot shows the gene effect score as calculated by DepMap (Arafeh et al. 2025) of SOX17 in cancer models from uterus, ovary/fallopian tube, breast, and all other lineages.

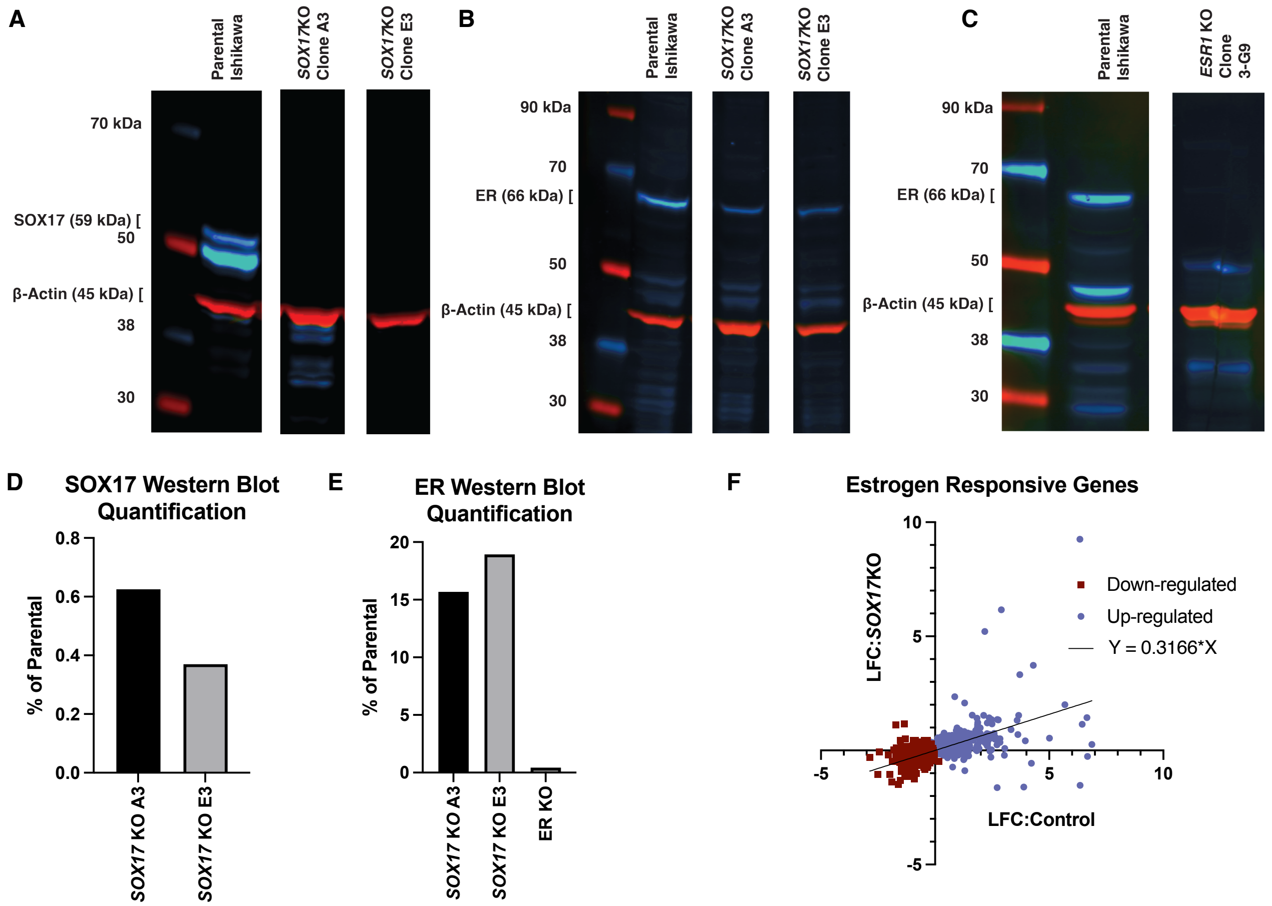


**Supplemental Figure 9: SOX17 and ER KO affect protein levels.** A) Western blot shows absence of SOX17 protein in SOX17 KO lines (SOX17 in blue, beta actin in red). B) Western blot shows reduced ER protein in SOX17 KO lines (ER in blue, beta actin in red). C) Western blot shows absence of ER protein in ER KO line (ER in blue, beta actin in red). Bar graphs show Western blot quantification for D) panel A and E) panels B and C. F) Scatter plot shows $log_2$(fold change) of estrogen responsive genes in SOX17 KO lines compared to control lines, with linear regression line shown in black. This figure is related to Figure 6.

### Supplemental Table 1: Datasets used in the Boruta analyses

| Cell Line | Factor | Reference | GEO Accession |
|---|---|---|---|
| Ishikawa | ATAC-seq | (Vahrenkamp et al. 2018) | GSE109893 |
| Ishikawa | CTCF | ENCODE (ENCODE Project Consortium 2012) | GSE32465 |
| Ishikawa | DNase-seq | ENCODE (ENCODE Project Consortium 2012) | GSE32970 |
| Ishikawa | ETV4 | ENCODE (ENCODE Project Consortium 2012) | GSE32465 |
| Ishikawa | FOXA1 | ENCODE (ENCODE Project Consortium 2012) | GSE32465 |
| Ishikawa | GR | (Vahrenkamp et al. 2018) | GSE109893 |
| Ishikawa | H3K27ac 1hr | (Vahrenkamp et al. 2018) | GSE109893 |
| Ishikawa | H3K27ac 8hr | (Vahrenkamp et al. 2018) | GSE109893 |
| Ishikawa | H3K4me1 | (Ginley-Hidinger et al. 2024) | GSE227241 |
| Ishikawa | H3K4me3 | (Ginley-Hidinger et al. 2024) | GSE227241 |
| Ishikawa | Hi-ChIP | (Ginley-Hidinger et al. 2024) | GSE227242 |
| Ishikawa | JUN | (Ginley-Hidinger et al. 2024) | GSE227241 |
| Ishikawa | LSD1 | (Ginley-Hidinger et al. 2024) | GSE227241 |
| Ishikawa | MAX | (Ginley-Hidinger et al. 2024) | GSE227241 |
| Ishikawa | MYC | (Ginley-Hidinger et al. 2024) | GSE227241 |
| Ishikawa | PRO-seq | (Ginley-Hidinger et al. 2024) | GSE227243 |
| Ishikawa | RARA | (Ginley-Hidinger et al. 2024) | GSE227241 |
| Ishikawa | Rel Affinity | This work | |
| Ishikawa | RNAPII | ENCODE (ENCODE Project Consortium 2012) | GSE32465 |
| Ishikawa | SIN3A | (Ginley-Hidinger et al. 2024) | GSE227241 |
| Ishikawa | SOX17 | This work | GSE 312323 |
| Ishikawa | TAF1 | (Ginley-Hidinger et al. 2024) | GSE227241 |
| T-47D | ATAC-seq | (Arnesen et al. 2021) | GSE148278 |
| T-47D | BRD4 | (Shu et al. 2016) | GSE63581 |
| T-47D | CTCF | (Gertz et al. 2013) | GSE32465 |
| T-47D | DNase-seq | ENCODE (ENCODE Project Consortium 2012) | GSE32970 |
| T-47D | FOXA1 | ENCODE (ENCODE Project Consortium 2012) | GSE32465 |
| T-47D | GATA3 | ENCODE (ENCODE Project Consortium 2012) | GSE32465 |
| T-47D | H3K27ac | (Carleton et al. 2020) | GSE147141 |
| T-47D | H3K27me3 | (Ginley-Hidinger et al. 2024) | GSE227241 |
| T-47D | H3K4me1 | (Ginley-Hidinger et al. 2024) | GSE227241 |
| T-47D | H3K4me3 | (Ginley-Hidinger et al. 2024) | GSE227241 |
| T-47D | Hi-ChIP | (Ginley-Hidinger et al. 2024) | GSE227242 |
| T-47D | JUN | (Ginley-Hidinger et al. 2024) | GSE227241 |
| T-47D | LSD1 | (Ginley-Hidinger et al. 2024) | GSE227241 |
| T-47D | MAX | (Ginley-Hidinger et al. 2024) | GSE227241 |
| T-47D | PRO-seq | (Ginley-Hidinger et al. 2024) | GSE227243 |
| T-47D | RARA | (Ginley-Hidinger et al. 2024) | GSE227241 |
| T-47D | ERE Rel Affinity | This work | |
| T-47D | RNAPII | (Ginley-Hidinger et al. 2024) | GSE227241 |

| | | | |
|---|---|---|---|
| T-47D | SIN3A | (Ginley-Hidinger et al. 2024) | GSE227241 |
| T-47D | TAF1 | (Ginley-Hidinger et al. 2024) | GSE227241 |

### Supplemental Table 6: Primer Sequences

| Primer Name | Sequence |
|---|---|
| 3XERE PCR Forward Primer | TAAGCAAAGCTTCGAGCTCTTACGCGTGCTA |
| 3XERE PCR Reverse Primer | TAAGCAATCGATCGAGCTCTTACGCGTGCTA |
| CRISPR Screen Genomic DNA PCR Amplification Forward Primer | TCGTCGGCAGCGTCAGATGTGTATAAGAGA CAGTTGTGGAAAGGACGAAACACCG |
| CRISPR Screen Genomic DNA PCR Amplification Reverse Primer | GTCTCGTGGGCTCGGAGATGTGTATAAGAG ACAGCCAATTCCCACTCCTTTCAAGACCT |
| Index 1 Read Adapter for CRISPR Screen | CAAGCAGAAGACGGCATACGAGAT[i7]GTC TCGTGGGCTCGG |
| Index 2 Read Adapter for CRISPR Screen | AATGATACGGCGACCACCGAGATCTACAC [i5]TCGTCGGCAGCGTC |
| SOX17 KO gRNA 1 Forward | CACCGGCCGAGTTGAGCAAGATGCT |
| SOX17 KO gRNA 1 Reverse | AAACAGCATCTTGCTCAACTCGGCC |
| SOX17 KO gRNA 2 Forward | CACCGAGGGCGAGTCCCGTATCCGG |
| SOX17 KO gRNA 2 Reverse | AAACCCGGATACGGGACTCGCCCTC |
| ER KO gRNA Forward | CACCGGGGATACGAAAAGACCGAAG |
| ER KO gRNA Reverse | AAACCTTCGGTCTTTTCGTATCCCC |
